# Two Type VI Secretion Systems in *Vibrio coralliilyticus* RE22Sm exhibit differential target specificity for bacteria prey and oyster larvae

**DOI:** 10.1101/2021.05.28.446209

**Authors:** Christian W. Schuttert, Marta Gomez-Chiarri, David C. Rowley, David R. Nelson

## Abstract

*Vibrio coralliilyticus* is an extracellular bacterial pathogen and a causative agent of vibriosis in larval oysters. Host mortality rates can quickly reach 100% during vibriosis outbreaks in oyster hatcheries. Type VI Secretion Systems (T6SS) are rapidly polymerizing, contact dependent injection apparatus for prey cell intoxication and play important roles in pathogenesis. DNA sequencing of *V. coralliilyticus* RE22Sm indicated the likely presence of two functional T6SSs with one on each of two chromosomes. Here, we investigated the antibacterial and anti-eukaryotic roles of the two T6SSs (T6SS1 and T6SS2) against *E. coli* Sm10 cells and *Crassostrea virginica* larvae, respectively. Mutations in *hcp* and *vgrG* genes were created and characterized for their effects upon bacterial antagonism and eukaryotic host virulence. Mutations in *hcp1* and *hcp2* resulted in significantly reduced antagonism against *E. coli* Sm10, with the *hcp2* mutation demonstrating the greater impact. In contrast, mutations in *vgrG1* or *vgrG2* had little effect on *E. coli* killing. In eastern oyster larval challenge assays, T6SS1 mutations in either *hcp1* or *vgrG1* dramatically attenuated virulence against *C. virginica* larvae. Strains with restored wild type *hcp* or *vgrG* genes reestablished T6SS-mediated killing to that of wild type *V. coralliilyticus* RE22Sm. These data suggest that the T6SS1 of *V. coralliilyticus* RE22Sm principally targets eukaryotes and secondarily bacteria, while the T6SS2 primarily targets bacterial cells and secondarily eukaryotes. Attenuation of pathogenicity was observed in all T6SS mutants, demonstrating the requirement for proper assembly of the T6SS systems to maintain maximal virulence.

**Importance:** Vibriosis outbreaks lead to large-scale hatchery losses of oyster larvae (product and seed) where *Vibrio* sp. associated losses of 80 to 100 percent are not uncommon. Practical and proactive biocontrol measures can be taken to help mitigate larval death by *Vibrio* sp. by better understanding the underlying mechanisms of virulence in *V. coralliilyticus*. In this study, we demonstrate the presence of two Type VI Secretion Systems (T6SS) in *V. coralliilyticus* RE22Sm and interrogate the roles of each T6SS in bacterial antagonism and pathogenesis against a eukaryotic host. Specifically, we show that the loss of T6SS1 function results in the loss of virulence against oyster larvae.

## INTRODUCTION

Bacterial diseases in aquatic environments negatively affect development and advancement of aquaculture systems throughout the world (1–3). *Vibrio* species are among the most common bacterial pathogens in marine aquaculture settings (4). *Vibrio coralliilyticus* RE22Sm (formerly classified as *V. tubiashii* RE22) is a Gram-negative motile marine bacterium and a member of the *Vibrionaceae* within the γ-proteobacteria class (5). *V. coralliilyticus* RE22Sm is a bacterial pathogen of larval eastern (*Crassostrea virginica*) and Pacific oysters (*Crassostrea gigas*) and has been associated with major disease outbreaks in hatcheries, causing shortages in seed oysters for commercial shellfish producers (1, 6). Mortality from *V. coralliilyticus* induced vibriosis can rapidly reach 100% in larval rearing tanks and contributes to significant economic losses to bivalve aquaculture worldwide (7). Historically, antimicrobial agents have been used to combat disease outbreaks in aquaculture. However, their usage is discouraged due to rapidly emerging antibiotic resistance and the toxicity of many of these agents (8–10). Improved knowledge on mechanisms of *Vibrio* pathogenicity in oysters would be useful in developing alternative disease management strategies.

Previous studies of virulence factors employed by *V. coralliilyticus* strains have mainly focused on extracellular enzyme function, as it was thought to be the driving force of pathogenicity (11). In addition to protease production, the annotated genome for RE22Sm (12) provides evidence for additional potential virulence factors, including a Type 3 Secretion System (T3SS), an RtxA-like toxin with its dedicated Type 1 Secretion System (T1SS), a pore-forming hemolysin (homologous to the Vah1 hemolysin of *Vibrio anguillarum*), a phospholipase hemolysin (homologous to the Plp hemolysin of *V. anguillarum*), several secreted proteases, and two Type VI Secretion Systems (T6SS) (13, 14).

The T6SS is a contact-mediated bacterial nanomachine composed of thirteen conserved proteins that inject effector proteins directly into a eukaryotic or bacterial cell (15). Many effector proteins translocated into the host/prey cell are bound as cargo to the proteins that constitute the physical T6SS puncturing device. This puncturing device is comprised of the hemolysin co-regulated protein (Hcp) and the valine glycine repeat protein G (VgrG). Specialized elongated versions of Hcp and VgrG that contain effector domains may also act as effectors (16, 17). The T6SS has been shown to be vital for virulence in many organisms, including *Vibrio cholera*e, where the T6SS was first discovered (18). Against eukaryotic prey, effector proteins can modify the host cell membrane to facilitate penetration, evade the phagosome, spread intracellularly, and cause direct cytotoxic effects (19). Moreover, the T6SS may enable Gram-negative bacteria to kill and out-compete other species of bacteria that occupy a similar niche (20). Effector proteins have been shown to cause complete lysis of other Gram-negative bacterial cells via membrane-targeting phospholipases, peptidoglycan-targeting amidases and glycoside hydrolases (21). Other antimicrobial effectors have DNase activity (22). Endogenous immunity proteins prevent a bacterium containing a T6SS from harming sibling cells. Some species of bacteria, such as *Pseudomonas aeruginosa* and *V. cholerae*, utilize their T6SS to translocate both antimicrobial and anti-eukaryotic effectors (23). In this report, we describe two T6SSs used by *V. coralliilyticus* RE22Sm as antibacterial and virulence factors. These findings provide new insights into the mechanisms by which RE22Sm eliminates bacterial competition and promotes pathogenesis in oyster larvae.

## RESULTS

### The RE22Sm genome contains two distinct T6SSs

Two distinct T6SS associated gene clusters were identified by utilizing a bioinformatics-guided approach to survey the annotated *V. coralliilyticus* RE22Sm genome (5, 12). Initially, genes were identified using Rapid Annotation using Subsystem Technology (RAST) (24). Twenty-two genes on chromosome 1 (Table S1) spanning 27,726 bp with a G+C ratio of 42% suggested the presence of a type VI secretion system (T6SS1). Twenty-seven genes on chromosome 2 (Table S2) spanning 25,060 bp with a G+C ratio of 43.1% were suggestive of a second system, T6SS2. The G+C content of both T6SS gene clusters was slightly lower than the entire RE22Sm genome (45.8%). Genes and motifs were identified and analyzed as previously described by Solomon *et al*. (23) to identify <u>m</u>arkers for type s<u>ix</u> effector (MIX) motifs (Table 1). Five MIX motifs were located including four within the T6SS2 gene cluster. Two MIX motifs were found in both TssA2/ImpA2 and TssI2/VgrG2. An additional MIX motif was located outside the T6SS2 gene cluster in a hypothetical protein identified as a possible oxalate:formate antiporter. The top 100 hits for this protein on BLASTx exhibited >92% amino acid sequence identity and all were in *Vibrio* species. No MIX motif-containing genes were found in the T6SS1 or elsewhere in the RE22Sm genome.

**Table 1.**
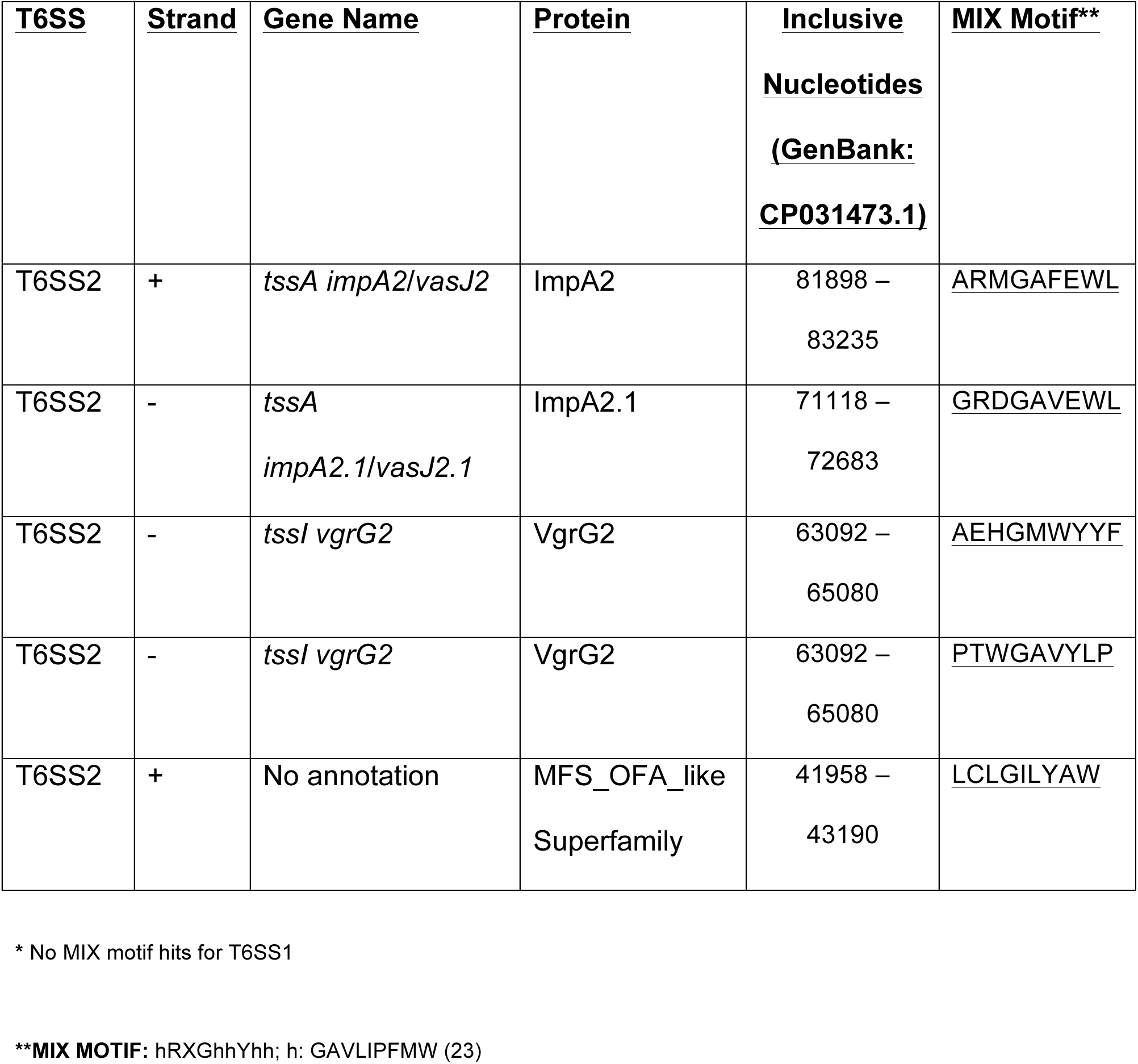
T6SS MIX motif search in V. coralliilyticus RE22Sm (12)

| <u>T6SS</u> | <u>Strand</u> | <u>Gene Name</u> | <u>Protein</u> | <u>Inclusive</u><br><u>Nucleotides</u><br><u>(GenBank:</u><br><u>CP031473.1)</u> | <u>MIX Motif**</u> |
| --- | --- | --- | --- | --- | --- |
| T6SS2 | + | *tssA impA2/vasJ2* | ImpA2 | 81898 –<br>83235 | <u>ARMGAFEWL</u> |
| T6SS2 | - | *tssA*<br>*impA2.1/vasJ2.1* | ImpA2.1 | 71118 –<br>72683 | <u>GRDGAVEWL</u> |
| T6SS2 | - | *tssI vgrG2* | VgrG2 | 63092 –<br>65080 | <u>AEHGMWYYF</u> |
| T6SS2 | - | *tssI vgrG2* | VgrG2 | 63092 –<br>65080 | <u>PTWGAVYLP</u> |
| T6SS2 | + | No annotation | MFS_OFA_like<br>Superfamily | 41958 –<br>43190 | <u>LCLGILYAW</u> |

\* No MIX motif hits for T6SS1
**\*\*MIX MOTIF:** hRXGhhYhh; h: GAVLIPFMW (23)

Genes for one hemolysin co-regulated protein (*hcp1* and *hcp2*) and one valine glycine repeat protein G (*vgrG1* and *vgrG2*) were detected in each T6SS (Tables S1 and S2). The amino acid sequences of Hcp1 and Hcp2 shared 24% identity (E-value 5e^-04^), while the amino acid sequences of VgrG1 and VgrG2 shared 30% identity (E-value 1e^-75^). VgrG1 shared 86% identity with the VgrG1 protein of *V. cholerae* serotype O1, which also contains a PAAR (proline, alanine, alanine, arginine) motif (25). The *V. cholerae* VgrG1 functions as an actin cross-linking toxin in eukaryotes and a toxic effector toward bacteria (18, 26, 27). The actin cross-linking domain (ACD) of *V. cholerae* is unique to this organism and was not detected in the RE22Sm VgrG1 protein. A PAAR motif is encoded in the small *paaR* gene downstream from *vgrG1* of RE22Sm (Table S1). No PAAR motif was detected in the VgrG2 of T6SS2 (Table S2), although a possible lysozyme domain was detected. Endopeptidase and lysozyme domains were detected within a single putative extracellular protein of the M23 endopeptidase family (Accession number: **CP031473.1**) in the T6SS2 gene cluster of both *V. coralliilyticus* RE22Sm and *V. coralliilyticus* BAA-450 (YB1) (99% identity to RE22Sm M23 containing protein). This 320 amino acid protein is not associated with any annotated gene or effector, but is detectable in other *Vibrio* species (12).

The previously reported *V. coralliilyticus* YB1 genome (28) was compared to the RE22Sm genome to help determine the organization of the RE22Sm T6SS. One hypothetical protein containing a Ras-GTPase activity domain (Accession number: **WP_006962091.1**) is encoded near the genes of the T6SS1 core machinery and may serve as an effector protein against host cells (28). Ras-GTPase activating domains enhance hydrolysis of GTP bound to Ras-GTPases. A mechanism of this type has yet to be demonstrated in T6SS effector compounds, but has been found in toxins secreted by T3SS in *Yersinia pseudotuberculosis* and *Pseudomonas aeruginosa* (15).

### Antibacterial T6SS activity of *V. coralliilyticus* RE22Sm against *E. coli* Sm10

We examined the antibacterial activity of the RE22Sm T6SSs by combining RE22Sm cells (attacking cells) and *E. coli* Sm10 (prey cells) on filters at a multiplicity of infection (MOI) = 4 for 4 h (Fig. 1) in a standard T6SS assay (as described in the Materials and Methods section). Incubation of RE22Sm with *E. coli* Sm10 consistently resulted in a >3 log decline the *E. coli* Sm10 cells. Knockout mutations in *hcp1*, *hcp2*, *vgrG1*, and *vgrG2* were constructed and the resulting mutant strains tested for their ability to kill *E. coli* Sm10 (Fig. 2). Knockout (KO) mutations in *hcp1* (Fig. 2a) or *hcp2* (Fig. 2b) significantly reduced killing of the target *E. coli* Sm10 cells by 2-3 orders of magnitude when compared to T = 0 h *E. coli* Sm10 CFU/ml. *E. coli* Sm10 cells declined by 1.2 log (*P* < 0.05) and 0.46 log (*P* < 0.01) when incubated with the *hcp1* and *hcp2* mutants for 4 h, respectively, as compared to a decline of 3.38 log when incubated with wild type RE22Sm cells. In contrast, mutations in either *vgrG1* (Fig. 2a) or *vgrG2* (Fig. 2b) had no significant effect on the viability of the target cells (*E. coli* declines of 3.58 log and 3.15 log, respectively). *In cis* (Fig. 2c) and *in trans* (Fig. 2d) complements of *hcp1*, *hcp2*, *vgrG1*, and *vgrG2* reversed the effects of the mutations demonstrating that knockouts of these T6SS genes affect prey killing. Further, when double KO mutants of *hcp1* and *hcp2* were tested, no significant killing of the *E. coli* target cells was detected (n.s.) (decline of 0.07 log or >85% survival). *E. coli* Sm10 cells declined by 1.2 log (*P* < 0.001) when incubated with the *hcp1/2* mutant for 4 h, compared to a decline of 3.38 log when incubated with wild type RE22Sm. Double KO mutants of *vgrG1* and *vgrG2* (Fig. 3) exhibited significantly impaired killing ability (*P* < 0.005) (decline of 1.68 log or 2.1% survival) when compared to the RE22Sm control (decline of 3.03 log). *E. coli* Sm10 cells incubated with *vgrG1/2* for 4 h declined 1.8 log (P < 0.05) when compared to RE22Sm wild type decline of 3.38 log.

**Figure 1.**
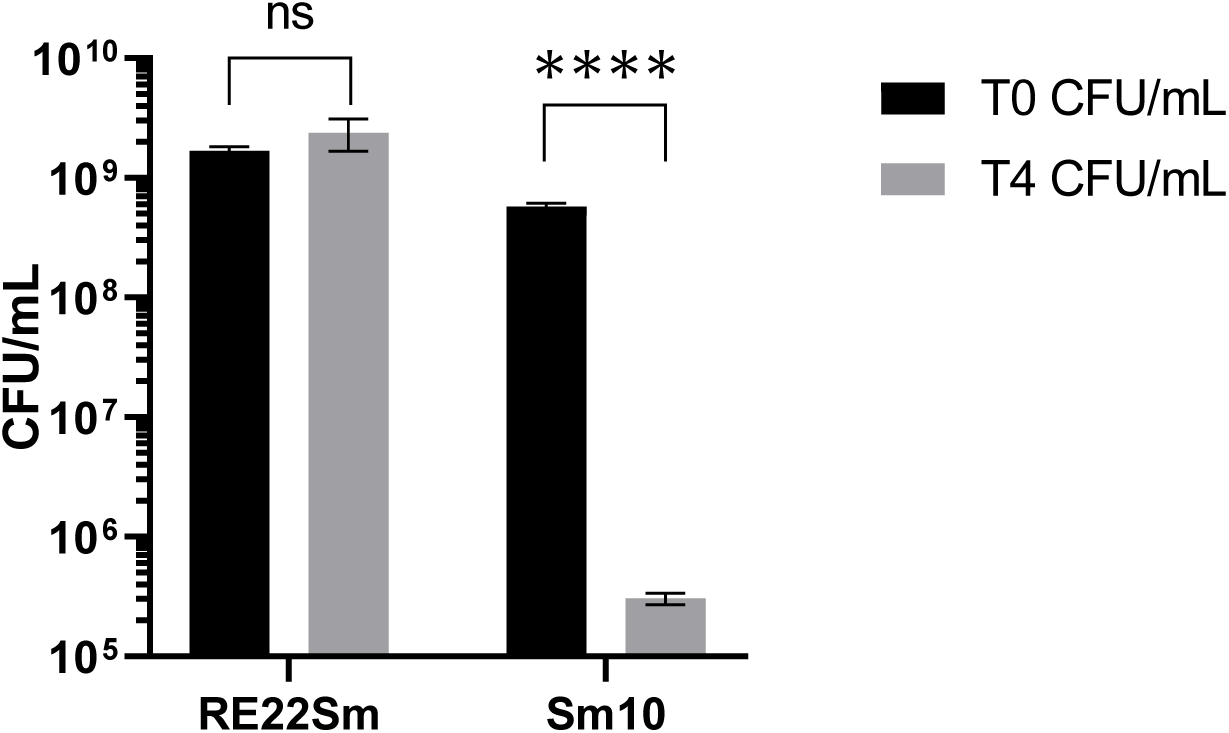
Determination of the *V. coralliilyticus* RE22Sm (attacking cell) T6SS-mediated antibacterial activity against *E. coli* Sm10 (prey cell) when incubated on a filter for 4 h at 27°C with a 4:1 predator: prey ratio (MOI = 4). Starting RE22 cell density was ∼2×10^9^ CFU/ml and starting *E. coli* Sm10 cell density was ∼5 ×10^8^ CFU/ml. The data are the average of 3 biological replicates (experiments); each experiment had three technical replicates. Error bars represent ±1 standard deviation (SD). Statistical analysis by Student’s T-test. ns = not significant, **** *= P* < 0.001

**Figure 2.**
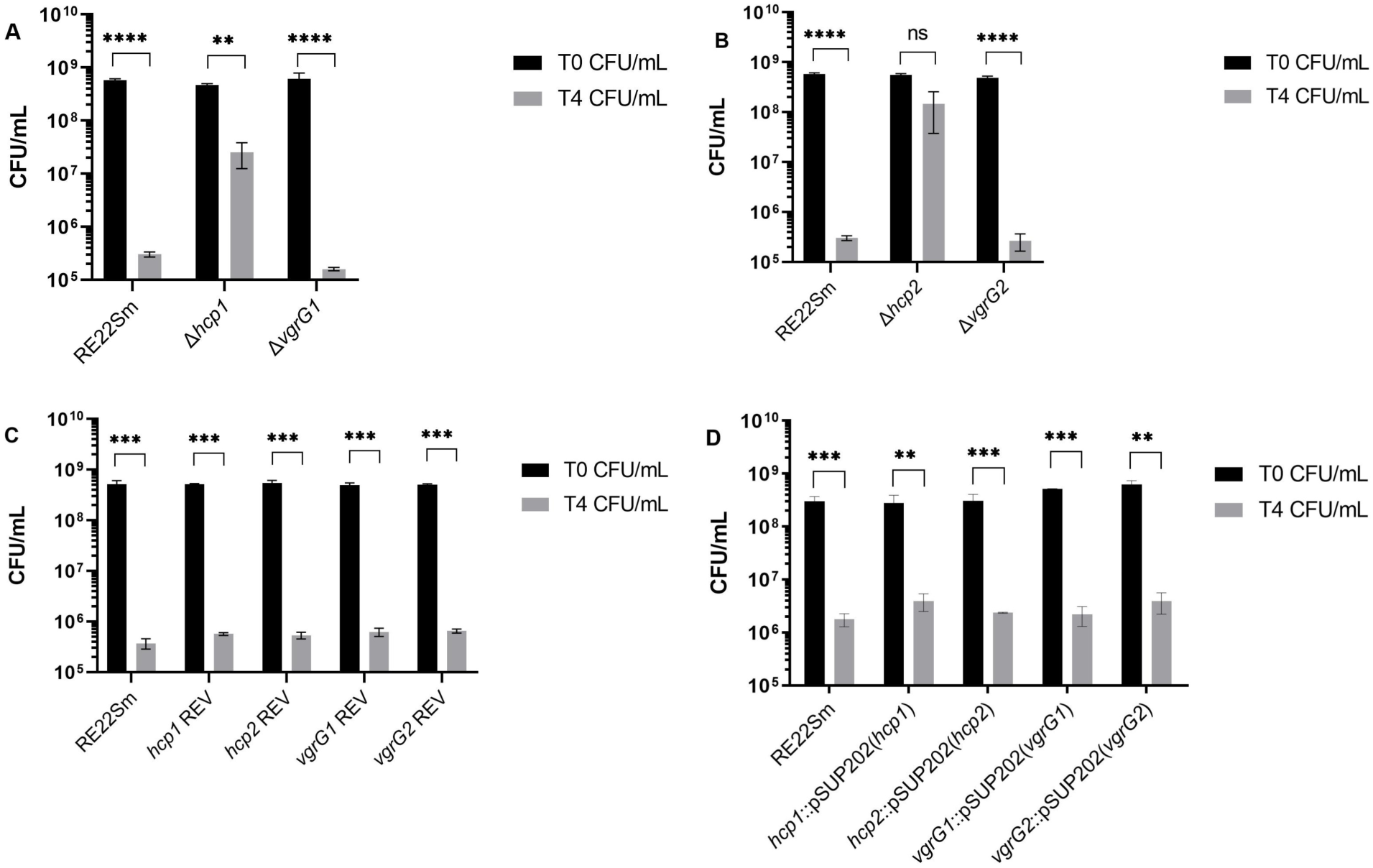
T6SS-mediated anti-bacterial activity of *V. coralliilyticus* RE22Sm wild type and T6SS mutant strains against *E. coli* Sm10 prey cells. Each group of two bars shows the cell density (CFU/ml) of the *E. coli* Sm10 prey cells at T= 0 h (black bar) and T= 4 h (grey bar) after being mixed with attacking *V. coralliilyticus* wild type (RE22Sm) or T6SS mutant strains. **(A)** T6SS killing activity of RE22Sm mutant strains Δ*hcp1*, Δ*vgrG1*, and RE22Sm wild-type control. **(B)** T6SS2 killing assay by RE22Sm mutant strains Δ*hcp2*, Δ*vgrG2*, and RE22Sm wild-type control. **(C)** T6SS killing assay by RE22Sm T6SS mutant revertant, strains and RE22Sm wild-type control. **(D)** T6SS killing assay by RE22Sm T6SS mutant *in-trans* complement strains, and RE22Sm wild-type control. All data are averages of at least 3 experiments; error bars show ±1 SD; ns = not significant, * = *P* < 0.05, ** = *P* < 0.01, *** = *P* < 0.005, **** *= P* < 0.001 (Statistical analysis by unpaired Student’s T-test).

**Figure 3.**
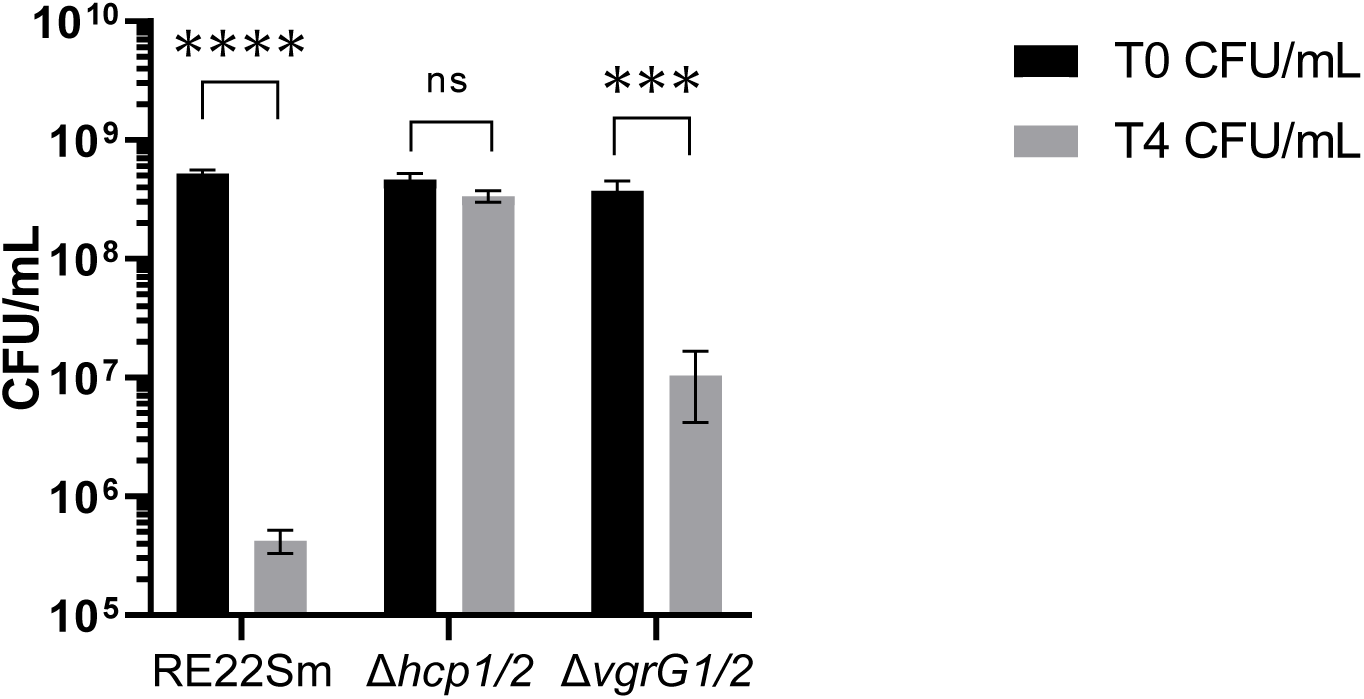
T6SS killing activity of *V. coralliilyticus* RE22Sm wild-type and T6SS double mutants against *E. coli* Sm10 prey cells. Each group of two bars shows the cell density (CFU/ml) of the *E. coli* Sm10 prey cells at T= 0 h (black bar) and T =4h (grey bar) after being mixed with attacking *V. coralliilyticus* wild type (RE22Sm) or T6SS mutant strains. Average of 3 experiments; error bars indicate ±1 SD; ns = not significant, * = *P* < 0.05, ** = *P* < 0.01, *** = *P* < 0.005, **** *= P* < 0.001 (Statistical analysis by unpaired Student’s T-test).

We also examined the possibility that other potential virulence-related genes might play a role in antibacterial activity. Allelic exchange mutations in the protease genes *vcpA* and *vcpB* and the transcriptional regulatory gene *vcpR* were constructed and the resulting strains of RE22Sm tested for their ability to kill *E. coli* target cells. The *vcpA* or the *vcpB* mutations had no effect on the killing of target cells (declines of 3.49 log and 3.08 log, respectively) as compared to the RE22Sm control (decline of 3.33 log). The *vcpR* mutant reduced the *E. coli* cell density by 2.52 log, 0.81 log less killing of *E. coli* Sm10 target cells as compared to the wild type RE22Sm cells (Table 2).

**Table 2.** Effects of *V. coralliilyticus* protease mutants on T6SS-mediated antibacterial activity.^1^.

| Strain Mixtures | T0 CFU/mL (±1 SD) | T4 CFU/mL (±1 SD) |
| --- | --- | --- |
| **Mixture 1** |  |  |
| *V. coralliilyticus* RE22Sm† | 1.83E+09 (±1.6E+08) | 1.03E+09 (±7.6E+08) |
| *E. coli* Sm10* | 5.17E+08 (±5E+07) | 2.41E+05 (±6.8E+04) |
| **Mixture 2** |  |  |
| *V. coralliilyticus* Δ*vcpA*† | 1.43E+09 (±4.5E+08) | 3.10E+09 (±1.5E+08) |
| *E. coli* Sm10* | 5.33E+08 (±1.82E+07) | 1.73E+05 (±4.39E+04) |
| **Mixture 3** |  |  |
| *V. coralliilyticus* Δ*vcpB*† | 1.30E+09 (±4.6E+08) | 5.33E+09 (±4.8E+08) |
| *E. coli* Sm10* | 6.00E+08 (±3.6E+07) | 5.00E+05 (±3.5E+04) |
| **Mixture 4** |  |  |
| *V. coralliilyticus* Δ*vcpR*† | 1.67E+09 (±8.9E+08) | 6.20E+08 (±1.37E+08) |
| *E. coli* Sm10* | 4.44E+08 (±3.17E+08) | 1.34E+06 (±5.87E+05) |

<sup>1</sup> Predator cells† were mixed in a ratio of 4:1 with prey cells\* as described in the Materials and Methods section. The data are the average of 3 biological replicates (experiments); each experiment had three technical replicates.

### Antibacterial T6SS activity of *V. coralliilyticus* RE22Sm against *Vibrio anguillarum* strains

The T6SS assay was next used to determine the ability of RE22Sm to kill *Vibrio anguillarum* NB10Sm and M93Sm (serotypes O1 and O2, respectively) (Fig. 4). Examination of each *V. anguillarum* genome revealed that NB10Sm contains T6SS elements, while M93Sm does not. With RE22Sm as the attacking cell and NB10Sm or M93Sm as prey, both strains of *V. anguillarum* exhibited sensitivity to predation by RE22Sm (Fig. 4a). NB10Sm cell density declined by 1.81 orders of magnitude from 4.1×10^8^ CFU/ml to 6.37×10^6^ CFU/ml (*P* < 0.005), while M93Sm CFU/ml dropped 1.32 orders of magnitude from 4.5×10^8^ CFU/ml to 2.13×10^7^ CFU/ml (*P* < 0.01). These results suggest a partial, strain specific susceptibility or immunity to *V. coralliilyticus* RE22Sm T6SS effectors as compared to *E. coli* Sm10.

**Figure 4.**
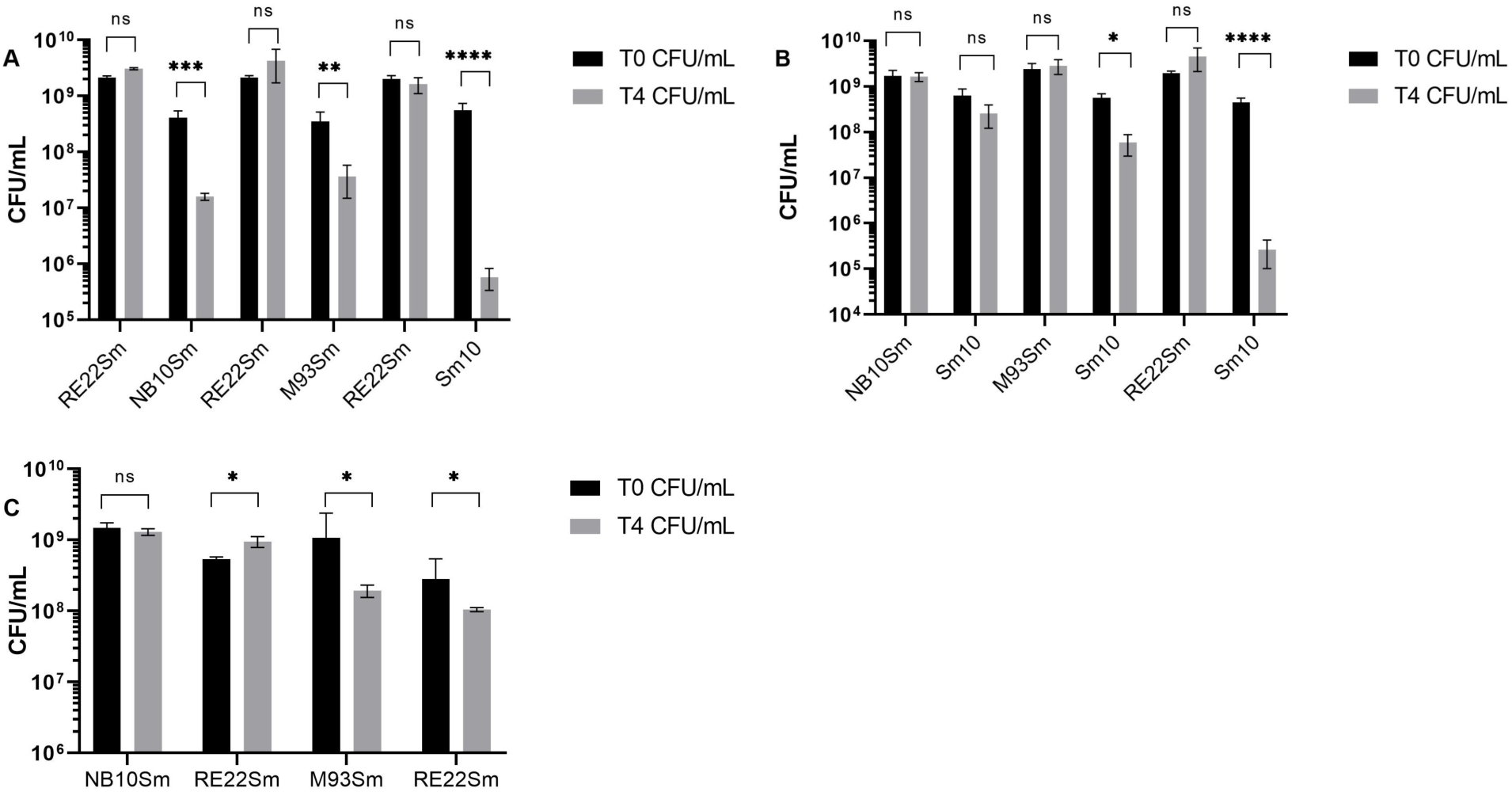
T6SS killing activity of *V. coralliilyticus* RE22Sm, *V. anguillarum* NB10Sm, and *V. anguillarum* M93Sm against *E. coli* Sm10 and *Vibrio* prey cells. Each group of four bars indicate attacker (first two bars) and prey cell (second two bars) cell density at T= 0 h (black bars) and 4 h (grey bars) **(A)** The ability of RE22Sm to kill serotype O1 (NB10Sm) and O2 (M93Sm) strains of *V. anguillarum*. **(B)** The ability of *V. anguillarum* NB10Sm and M93Sm to kill *E. coli* Sm10. **(C)** The ability of *V. anguillarum* NB10Sm and M93Sm to attack *V. coralliilyticus* RE22Sm. The data are the averages of at least 3 experiments; the error bars indicate ±1 SD; ns = not significant, * = *P* < 0.05, ** = *P* < 0.01, *** = *P* < 0.005, **** *= P* < 0.001 (Statistical analysis by unpaired Student’s T-test).

We also tested the ability of both *V. anguillarum* strains to kill *E. coli* Sm10 using the standard T6SS killing assay (Fig. 4b). Neither strain demonstrated virulence toward *E. coli* Sm10 comparable to that of *V. coralliilyticus* RE22Sm. Cell density of Sm10 declined by 50-60% when incubated with NB10 (n.s.), while incubation with M93Sm resulted in a ∼1 log decline in Sm10 viability (*P* < 0.05).

The effect of *V. anguillarum* strains NB10Sm and M93Sm on *V. coralliilyticus* RE22Sm viability was further examined when mixed at a ratio of 4:1 (Fig. 4c). In the presence of NB10Sm (∼2×10^9^ CFU/ml), RE22Sm cell density increased ∼1.8-fold from 5.4×10^8^ CFU/ml to 9.52×10^8^ CFU/ml over 4 h (*P* < 0.05) while the NB10Sm cell density of did not significantly change. In the presence of M93Sm, the RE22Sm cell density declined 2.7-fold, from 2.84×10^8^ CFU/ml to 1.05×10^8^ CFU/ml, over 4 h (*P* < 0.05). Interestingly, the M93Sm density declined 5.46-fold, from 1.06×10^9^ CFU/ml to 1.94×10^8^ CFU/ml (*P* < 0.05).

### T6SS contributes to virulence against *Crassostrea virginica* larvae

The contribution of the two T6SSs found in RE22Sm to oyster larval disease was evaluated by examining the effects of mutations in the *hcp* and *vgrG* genes. Wild type and mutant strains of RE22Sm were evaluated for their ability to kill larval oysters as described by Karim *et al.* (29). Oyster larvae infected with wild-type RE22Sm (positive infection control) exhibited 48% survival. In contrast, larvae infected with the Δ*hcp1* mutant (*P* < 0.001) or the Δ*vgrG1* mutant (*P* < 0.001) were significantly attenuated in killing compared to RE22Sm wild-type control, and were not significantly different from the no treatment control (Fig. 5). Larvae infected with the Δ*hcp2* mutant (*P* < 0.005) or the Δ*vgrG2* mutant (*P* < 0.005) showed ∼74% and ∼84% survival, respectively.

**Figure 5:**
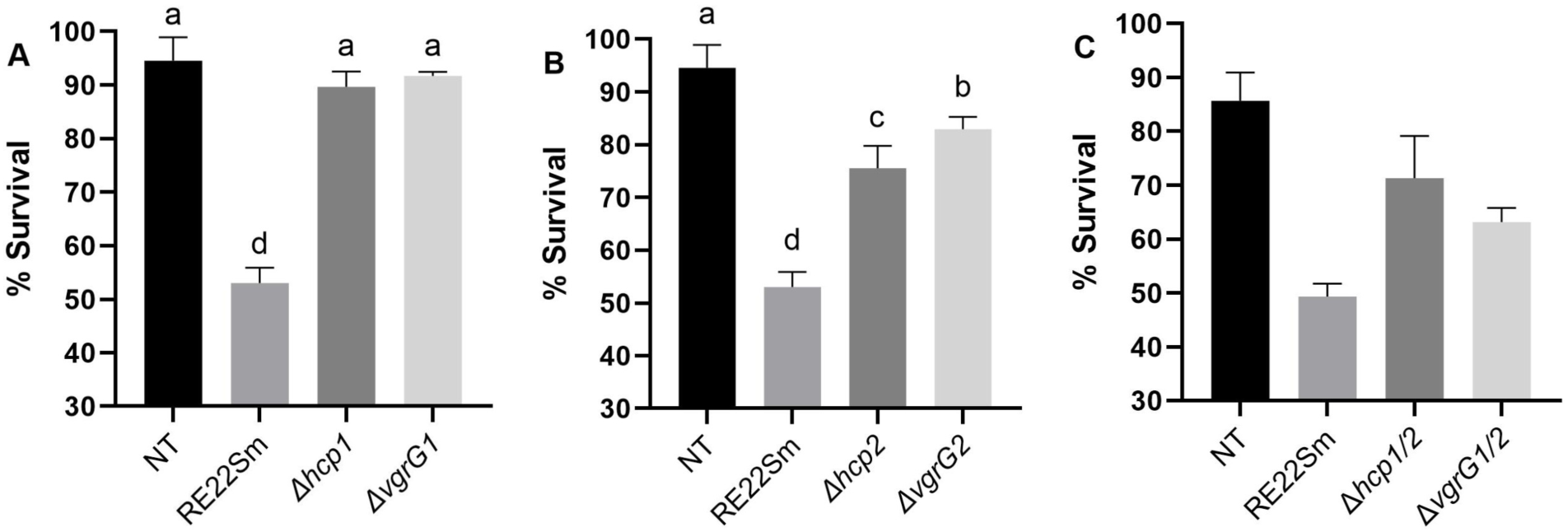
Oyster larvae survival after challenge *with V. coralliilyticus* RE22Sm. Oyster larvae were exposed to RE22Sm wild type and mutant strains (1×10^5^ CFU/ml) for 24 h. Oyster larvae treated with artificial seawater served as the negative control. Larval survival (% ±1 SD) was determined after 24 h challenge. **(A)** *V. coralliilyticus* RE22Sm wild type and T6SS1 mutants (Δ*vgrG1* and Δ*hcp1*) tested for virulence against larval oysters. **(B)** *V. coralliilyticus* RE22Sm wild type and T6SS2 mutants (Δ*vgrG2* and Δ*hcp2*) tested for virulence against larval oysters. **(C)** *V. coralliilyticus* RE22Sm wild type and T6SS double mutants *Δhcp1/2* and *ΔvgrG1/2* tested for virulence against larval oysters. Average of at least 3 biological replicates; the error bars indicate ±1 SD; different letters indicate statistical between among groups, a = *P* > 0.05, b = *P* < 0.05, c = *P* < 0.01, d = *P* < 0.005 (Statistical analysis by unpaired Student’s T-test, *P* < 0.05).

In comparison, the effects of KO mutations in the two proteases, *vcpA* and *vcpB*, and their transcriptional regulator, *vcpR*, previously identified as virulence factors in oyster pathogenesis (30), were examined. Eastern oyster larvae infected with the Δ*vcpA* (*P* < 0.005), Δ*vcpB* (*P* < 0.005), or Δ*vcpR* (*P* < 0.001) mutants survived at ∼71%, 72%, and 80%, respectively (Table S3).

Taken together, these results strongly suggest that T6SS1 is essential to RE22Sm virulence against larval oysters and that T6SS2, the VcpA and VcpB proteases, and the VcpR transcriptional regulator are important to successful pathogenesis and affect virulence, but are not essential.

## DISCUSSION

T6SSs are present in many Gram-negative bacteria and provide a means for bacterial competition and pathogenesis of eukaryotes (20). The soil pathogen *Burkholderia thailandensis* contains five distinct T6SSs that encompass a range of specificities toward different cell types (31). *V. cholerae* contains a single T6SS with dual function towards bacterial and eukaryotic target cells (32). Guillemette *et al.* (14) demonstrated that a functional T6SS in *V. coralliilyticus* OCN008 was necessary to kill strains of *V. cholerae*, adding to the repertoire of T6SSs identified in *Vibrio* species (33). A proteomic analysis of *V. coralliilyticus* YB1 supernatant detected sixteen T6SS proteins – all regulatory or structural in function (28).

In this study, we present data that *V. coralliilyticus* RE22Sm produces two functionally distinct T6SSs that act as virulence factors enabling these bacteria to attack both bacterial and eukaryotic targets. Analogous to Guillemette *et al*. (14), our RE22Sm strain can also kill the related *Vibrio* species, *V. anguillarum.* Initial detection of potential T6SS genes in RE22Sm utilized genomic findings in *V. coralliilyticus* YB1 (28) (Table S4). When the complete RE22Sm T6SS1 and T6SS2 were compared to the *Vibrionaceae* family by tBLASTx, all translated proteins were highly conserved throughout. Further, a comparative genomic approach assessing *V. coralliilyticus* virulence against *C. virginica* larvae indicates that the role of T6SS varies by bacterial strain and host/prey (34). Our results indicate that T6SS is required for pathogenicity and antibacterial activity in RE22Sm. Additionally, the multifaceted nature of the two T6SSs in *V. coralliilyticus* RE22Sm may allow their use in multiple steps during infection of oysters, such as clearing commensal bacteria, modification or killing of oyster cells and escaping the phagosome to allow intracellular spread within the host (35, 36).

Salomon *et al*. (23) proposed that some T6SS effectors in *Vibrio parahaemolyticus* could be identified by the presence of MIX motifs. We applied this idea to our inspection of the T6SSs of *V. coralliilyticus* RE22Sm. Despite the high degree of conservation of T6SSs across the *Vibrionaceae*, MIX motifs are not readily detected in RE22Sm by Find Individual Motif Occurrences (FIMO)(37). Further, we know that antibacterial activity is unaffected in the Δ*vgrG2* mutant, which contains two MIX motifs (Fig. 2b), and oyster virulence is only slightly attenuated (Fig. 5b). Consequently, the presence of MIX effectors is not required for T6SS activity, but is suggested to increase T6SS efficiency (23, 38).

Our data raised the question as to the roles of the two T6SSs in *V. coralliilyticus* RE22Sm antibacterial activity. We demonstrated that a deletion of either *hcp1* or *hcp2* results in a significant decline in the ability of these RE22Sm mutant strains to kill *E. coli* Sm10 prey cells when compared to the RE22Sm wild type. Complementation of either *hcp* gene (*cis* or *trans* complementation) restored predation activity to wild type levels. Further, knockouts of both *hcp1* and *hcp2* resulted in a near complete loss of bacterial killing. These data show that while both Hcp proteins are necessary for fully functional T6SS-mediated antibacterial activity, the loss of Hcp2 has a significantly larger effect upon predation. In contrast, deletion of either *vgrG1* or *vgrG2* has no significant effect upon predation compared to the RE22Sm wild type (Fig. 2). However, a double mutant for both *vgrG1* and *vgrG2* shows a significant decline in bacterial killing compared to the wild type RE22Sm. These data suggest that VgrG proteins contribute to predation, but only one of the two proteins is necessary for full activity. Therefore, both T6SS1 and T6SS2 possess antibacterial activity, with the loss of a functional Hcp2 having a having a somewhat larger effect on antibacterial activity than the loss of Hcp1.

VgrG switching, as described in *Serratia marcescens,* may account for the retention of function despite loss of either VgrG1 or VgrG2 is (39). Such switching capacity would allow the loaded VgrG, acting as an effector, to display preferential target specificity and the puncturing apparatus to be loaded according to the target organism (22). A second possibility is that only one complete T6SS system is necessary for antibacterial activity; however, the loss of either Hcp1 or Hcp2 has a much larger effect than the loss of either VgrG1 or VgrG2.

Guillemette *et al*. (14) examined the question of whether deletion mutations of protease genes *vtpA*, *vtpB* (renamed *vcpA* and *vcpB*) or their transcriptional regulator *vtpR* (*vcpR*) provide protection against predation by *V. cholerae* or affected T6SS-mediated killing of *V. cholerae* by *V. coralliilyticus* OCN008. They found that knockouts of *vcpA* and/or *vcpB* had no effect upon survival against killing by *V. cholerae* or ability to kill *V. cholerae*. However, the *vcpR* mutant had reduced ability to survive attack by *V. cholerae* and lost the ability to kill *V. cholerae*. We also found that deletion of either *vcpA* or *vcpB* in *V. coralliilyticus* RE22Sm had no effect on T6SS-mediated killing of prey cells. However, deletion of *vcpR* produced modest effect on predation of *E. coli* Sm10. The KO mutation of *vcpR* reduced antibacterial activity with target cell decline of 2.52 log compared to wild type RE22Sm causing *E. coli* Sm10 cell density to decline by 3.33 log, a reduction of ∼0.8 log. While we do not know the reason for the difference between effects of the *vcpR* mutation on predation, we suggest that *E. coli* Sm10 is a more vulnerable prey target than *V. cholerae*, perhaps because *V. cholerae* contains a T6SS with immunity genes (40) and *E. coli* Sm10 does not (41).

Our data also indicate that *V. coralliilyticus* RE22Sm is able to kill *V. anguillarum* strains NB10Sm and M93Sm (serotypes O1 and O2, respectively). However, both strains are significantly less sensitive to T6SS than *E. coli* Sm10. The decreased sensitivity of *V. anguillarum* to *V. coralliilyticus* T6SS-mediated predation may be due to the presence of immunity genes in their T6SS gene clusters. Tang *et al*. (2016) showed that *V. anguillarum* strains possess T6SS and are able to kill *E. coli* and *Edwardsiella tarda* (42). Our data demonstrate that NB10Sm is unable to kill either *E. coli* Sm10 (Fig. 4b) or *V. coralliilyticus* RE22Sm (Fig. 4c) in the T6SS assay, despite containing T6SS elements. However, while a search of the *V. anguillarum* M93Sm genome failed to reveal any T6SS genes, this O2 serotype strain is able to kill *E. coli* Sm10 in our T6SS assay. These results are of interest due to our initial hypothesis indicating that NB10Sm would be more virulent against *E. coli* Sm10 and RE22Sm than M93Sm due to the presence of T6SS genes in NB10Sm. We suggest that the T6SS of NB10Sm is inactive and that M93Sm has an unknown mechanism of antibacterial activity.

Our data begin to address the major role of the two T6SSs in *V. coralliilyticus* RE22Sm virulence against oysters. The hardened tip motif, PAAR, is present in the *paaR* protein found in T6SS1, possibly to puncture a thicker eukaryotic cell envelope (26) allowing a wider range of potential targets, including coral, the namesake target of *V. coralliilyticus*, and possibly other eukaryotes. This idea is supported by our observation that mutants lacking either *hcp1* or *vgrG1* are completely avirulent against oyster larvae, indicating that T6SS1 is required for pathogenesis against oyster larvae. In contrast, knockouts of either *hcp2* or *vgrG2* exhibited only partially attenuated virulence, suggesting that the T6SS2 plays a more limited role in pathogenesis of oyster larvae. A similar effect on virulence has been previously reported in *P. aeruginosa*, a microbe with multiple T6SSs under the transcriptional control of RpoN (σ^54^) (43). Further, contrary to expectations, RE22Sm mutants containing knockouts of both *hcp1* and *hcp2* or *vgrG1* and *vgrG2* were able to kill oyster larvae at greater rates than any of the single mutants in these genes. Understanding this observation will require further investigation, but does raise the possibility that other virulence genes are up-regulated when both T6SSs are knocked out.

The activities of the RE22Sm T6SSs together with other previously described virulence factors help to decode the pathogenic potential of this organism and demonstrate how this fast growing, motile organism can cause substantial mortality in an aquaculture setting. Increased understanding of *V. coralliilyticus* virulence genes involved in oyster infection should help inform efforts to prevent larval and juvenile vibriosis.

## MATERIALS & METHODS

### Bacterial strains, plasmids and growth conditions

*V. coralliilyticus* RE22Sm strains (Table 3) were routinely cultured in yeast peptone broth plus 3% NaCl (YP30), yeast peptone broth plus 3% Instant Ocean^©^ sea salt (mYP30), or Marine Minimal Medium (3M) plus 5% sucrose (44), supplemented with the appropriate antibiotic(s) in a shaking water bath (200 RPM) at 27°C. Overnight cultures of *V. coralliilyticus* RE22Sm, grown in mYP30, were harvested by centrifugation (8,000 × *g*; 10 min; 4°C), and the pelleted cells washed twice with sterile Nine Salt Solution (NSS) (45). Washed cells were resuspended to the appropriate cell densities in experimental media. *E. coli* strains were routinely cultured in LB20 (21). Antibiotics were used at the following concentrations: streptomycin, 200 µg/ml (Sm^200^); chloramphenicol, 5 µg/ml (Cm^5^) for *V. coralliilyticus*, and chloramphenicol, 20 µg/ml (Cm^20^) for *E. coli*; kanamycin, 50 µg/ml (Km^50^) for *E. coli*, kanamycin, 80 µg/ml (Km^80^) for *V. coralliilyticus* grown in liquid media, and kanamycin 80 µg/ml (Km^80^) for *V. coralliilyticus* grown on solid media. Agar plates were prepared using Difco Bacto^©^ agar at 1.6%.

**Table 3.** Bacterial strains and plasmids used in this study.

| Strain | Description | Resistance | Reference |
| --- | --- | --- | --- |
| ***V. coralliilyticus*** |  |  |  |
| RE22 | Wild-type isolate from oyster larvae |  | Estes *et al.* 2004 |
| RE22Sm | Spontaneous Sm<sup>r</sup> mutant of RE22 | Sm<sup>r</sup> | Zhao *et al.* 2016 |
| RE22Sm-GFP | Sm<sup>r</sup> Cm<sup>r</sup>; RE22Sm (pRhokHi-2-*gfp*) | Sm<sup>r</sup> Cm<sup>r</sup> | Zhao *et al.* 2016 |
| RE22SmKm | Sm<sup>r</sup> Km<sup>r</sup> mutant of RE22 harboring an empty pSUP203 shuttle vector | Sm<sup>r</sup> Km<sup>r</sup> | This study |
| RE22Δ*hcp1* | Sm<sup>r</sup> Km<sup>r</sup>; allelic exchange mutation of *hcp1* using pDM5; T6SS<sup>-/-</sup> | Sm<sup>r</sup> Km<sup>r</sup> | This study |
| RE22Δ*hcp2* | Sm<sup>r</sup> Km<sup>r</sup>; allelic exchange mutation of *hcp2* using pDM5; T6SS<sup>-/-</sup> | Sm<sup>r</sup> Km<sup>r</sup> | This study |
| RE22Δ*vgrG1* | Sm<sup>r</sup> Km<sup>r</sup>; allelic exchange mutation of *vgrG1* using pDM5; T6SS<sup>-/-</sup> | Sm<sup>r</sup> Km<sup>r</sup> | This study |
| RE22Δ*vgrG2* | Sm<sup>r</sup> Km<sup>r</sup>; allelic exchange mutation of *vgrG2* using pDM5; T6SS<sup>-/-</sup> | Sm<sup>r</sup> Km<sup>r</sup> | This study |
| RE22 Δ*hcp1*Δ*hcp2* | Sm<sup>r</sup> Cm<sup>r</sup>; Allelic exchange mutation of *hcp1* and *hcp2* using pDM5; T6SS<sup>-/-</sup> | Sm<sup>r</sup> Cm<sup>r</sup><br>Km<sup>r</sup> | This study |
| RE22<br>*ΔvgrG1ΔvgrG2* | Sm<sup>r</sup> Cm<sup>r</sup>; Allelic exchange mutation of *vgrG1* and *vgrG2* using pDM5; T6SS<sup>-/-</sup> | Sm<sup>r</sup> Cm<sup>r</sup><br>Km<sup>r</sup> | This study |
| RE22 *ΔvcpA* | Sm<sup>r</sup> Km<sup>r</sup>; allelic exchange mutation of *vcpA* using pDM4 | Sm<sup>r</sup> Cm<sup>r</sup> | This study |
| RE22 *ΔvcpB* | Sm<sup>r</sup> Km<sup>r</sup>; allelic exchange mutation of *vcpB* using pDM4 | Sm<sup>r</sup> Cm<sup>r</sup> | This study |
| RE22 *ΔvcpR* | Sm<sup>r</sup> Km<sup>r</sup>; allelic exchange mutation of *vcpR* using pDM4 | Sm<sup>r</sup> Cm<sup>r</sup> | This study |
| ***V. anguillarum*** |  |  |  |
| NB10SmKm | Spontaneous Sm<sup>r</sup> mutant of strain NB10 harboring an empty pSUP203 shuttle vector | Sm<sup>r</sup> Km<sup>r</sup> | This study |
| M93SmKm | Spontaneous Sm<sup>r</sup> mutant of strain M93 harboring an empty pSUP203 shuttle vector (contains Km resistance gene) | Sm<sup>r</sup> Km<sup>r</sup> | This study |
| ***E. coli*** |  |  |  |
| Sm10 | *Thi thr leu tonA lacY supE recA*<br>RP4-2 Tc::Mu::Km (λ) | Km<sup>r</sup> | Simon *et al.*,<br>1983 |
| Sm100 | Sm10 harboring pDM5 plasmid | Km<sup>r</sup> Cm<sup>r</sup> | This study |
| S122 | Sm10 harboring pSUP202P-<br>*gfp*(ORF) | Km<sup>r</sup> | Zhao *et al.* 2016 |
| CS01 | Sm10 harboring pDM5-*hcp1* |  |  |
| CS02 | Sm10 harboring pDM5-*hcp2* | Km<sup>r</sup> Cm<sup>r</sup> | This study |
| CS03 | Sm10 harboring pDM5-*vgrG1* | Km<sup>r</sup> Cm<sup>r</sup> | This study |
| CS04 | Sm10 harboring pDM5-*vgrG2* | Km<sup>r</sup> Cm<sup>r</sup> | This study |
| CS05 | Sm10 harboring pDM4-*vcpA* | Cm<sup>r</sup> | This study |
| CS06 | Sm10 harboring pDM4-*vcpB* | Cm<sup>r</sup> | This study |
| CS07 | Sm10 harboring pDM4-*vcpR* | Cm<sup>r</sup> | This study |
| **Plasmids** |  |  |  |
| pDM4 | Cm<sup>r</sup>; suicide vector with R6K origin and *sacB* | Cm<sup>r</sup> | Milton, 1996 |
| pDM5 | Cm<sup>r</sup> Km<sup>r</sup>; suicide vector with R6K origin and *sacB* | Cm<sup>r</sup> Km<sup>r</sup> | This study |
| pSUP202P | Ap<sup>r</sup> Cm<sup>r</sup> Tc<sup>r</sup>; broad host shuttle vector | Ap<sup>r</sup> Cm<sup>r</sup> Tc<sup>r</sup> | Simon *et al.*, 1983 |
| pSUP203 | Ap<sup>r</sup> Cm<sup>r</sup> Tc<sup>r</sup> Km<sup>r</sup>; broad host shuttle vector | Ap<sup>r</sup> Cm<sup>r</sup> Tc<sup>r</sup><br>Km<sup>r</sup> | This study |
| pRhokHi-2-*gfp* | pRhokHi-2-FbFP with *gfp* under the control of *PaphII* | Cm<sup>r</sup> | Zhao *et al.* 2016 |

### *V. coralliilyticus* RE22Sm bioinformatic analysis

*V. coralliilyticus* RE22Sm draft genome (LGLS00000000) was annotated by the RAST service (http://rast.nmpdr.org/rast.cgi) with default settings (24). A list of core genes and accessory components was compiled using T6SS information from *Pantoea ananatis* (46, 47), *Edwardsiella tarda* (48), and *Vibrio cholerae* (49–51). The MIX motif used was based on the findings of Salomon *et al*. (23). RE22Sm MIX motifs were detected by The MEME Suite – Find Individual Motif Occurrences (FIMO)(37) option, using default settings.

### Allelic exchange mutagenesis

The modified pDM4 plasmid containing a kanamycin resistance (Km^R^) gene, pDM5, was used to construct the allelic exchange mutants (Table 3) as described by Gibson *et al.* (52). The Km resistance gene was amplified from the TOPO2.1 vector (Invitrogen) and inserted into pDM5 via the Gibson Assembly Reaction at the AgeI restriction site. pDM5 was linearized at the SacI restriction enzyme site, using SacI-HF (New England Biolabs), within the multicloning region (MCR) for all mutation destined Gibson Assemblies. The ligation mixture was introduced into *E. coli* Sm10 (containing λpir) by electroporation with the BioRad Gene Pulser II in a 2 mm cuvette (2.5 kV; 25 µF; 200 Ω). Transformants were selected by growth on LB20Cm_20_ agar plates, and successful mutagenesis was confirmed by PCR screening for a novel junction between the pDM4 plasmid and the Gibson Fragment(s) from *V. coralliilyticus*. The mobilizable suicide vector was transferred from *E. coli* Sm10 into *V. coralliilyticus* RE22Sm by conjugation as previously described (53). Transconjugants were selected by utilizing the kanamycin resistance (Km^R^) gene located on the suicide plasmid. The subsequent incorporation of the target gene fragments into the suicide vector was confirmed by PCR analysis using specific primers (Table 4) to screen for the novel genetic inserts into the plasmid. The double crossover transconjugants were selected for by growth on 3MSm^200^ +5% sucrose agar plates for a second crossover event. Sucrose is used as the counter selective agent because pDM5 contains the *sacB* gene, which encodes levansucrase that converts sucrose to toxic levan (54). Putative allelic exchange mutants were screened for kanamycin sensitivity. The resulting RE22Sm mutants were then screened for the desired allelic exchange double crossover using PCR amplification.

**Table 4.**
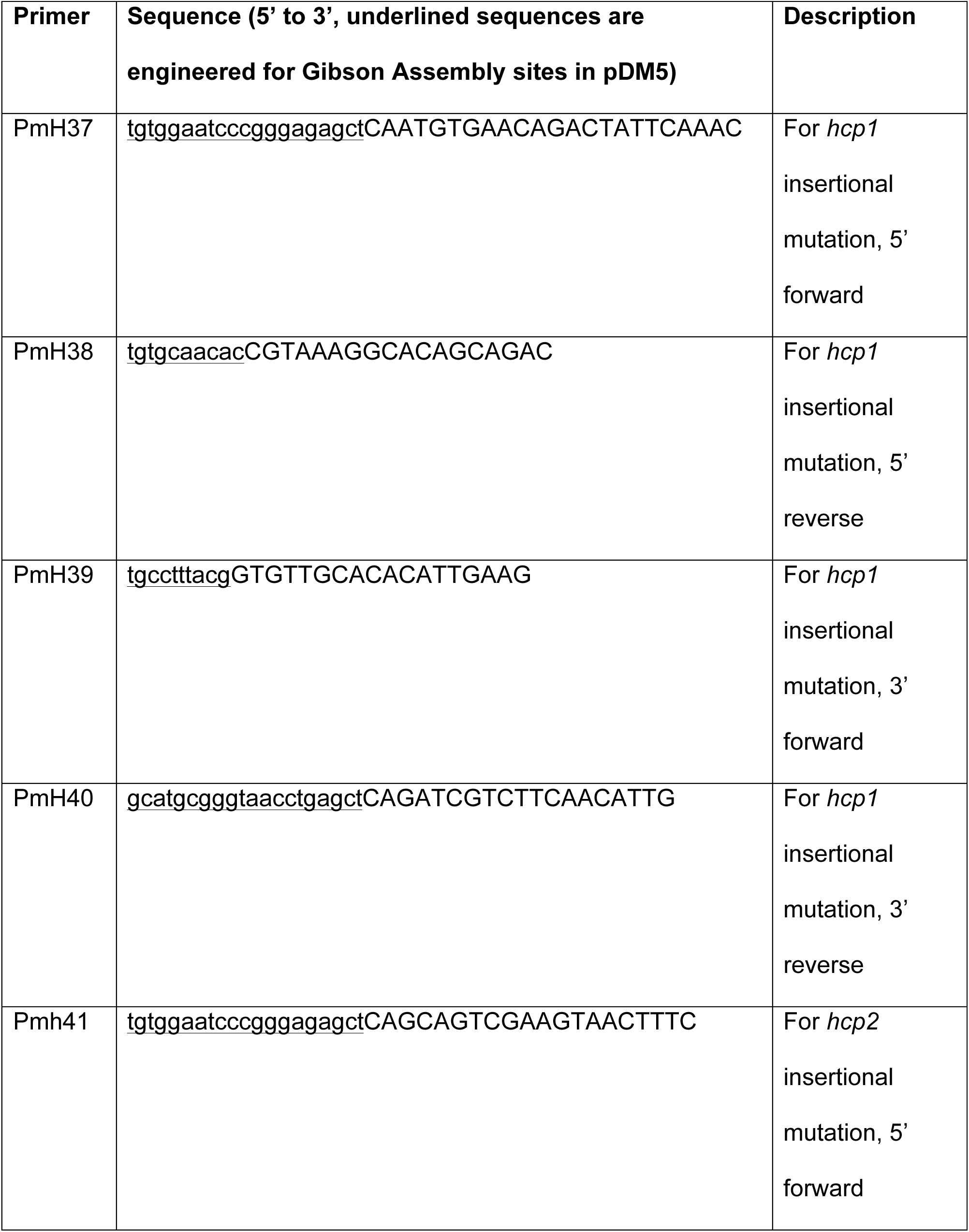
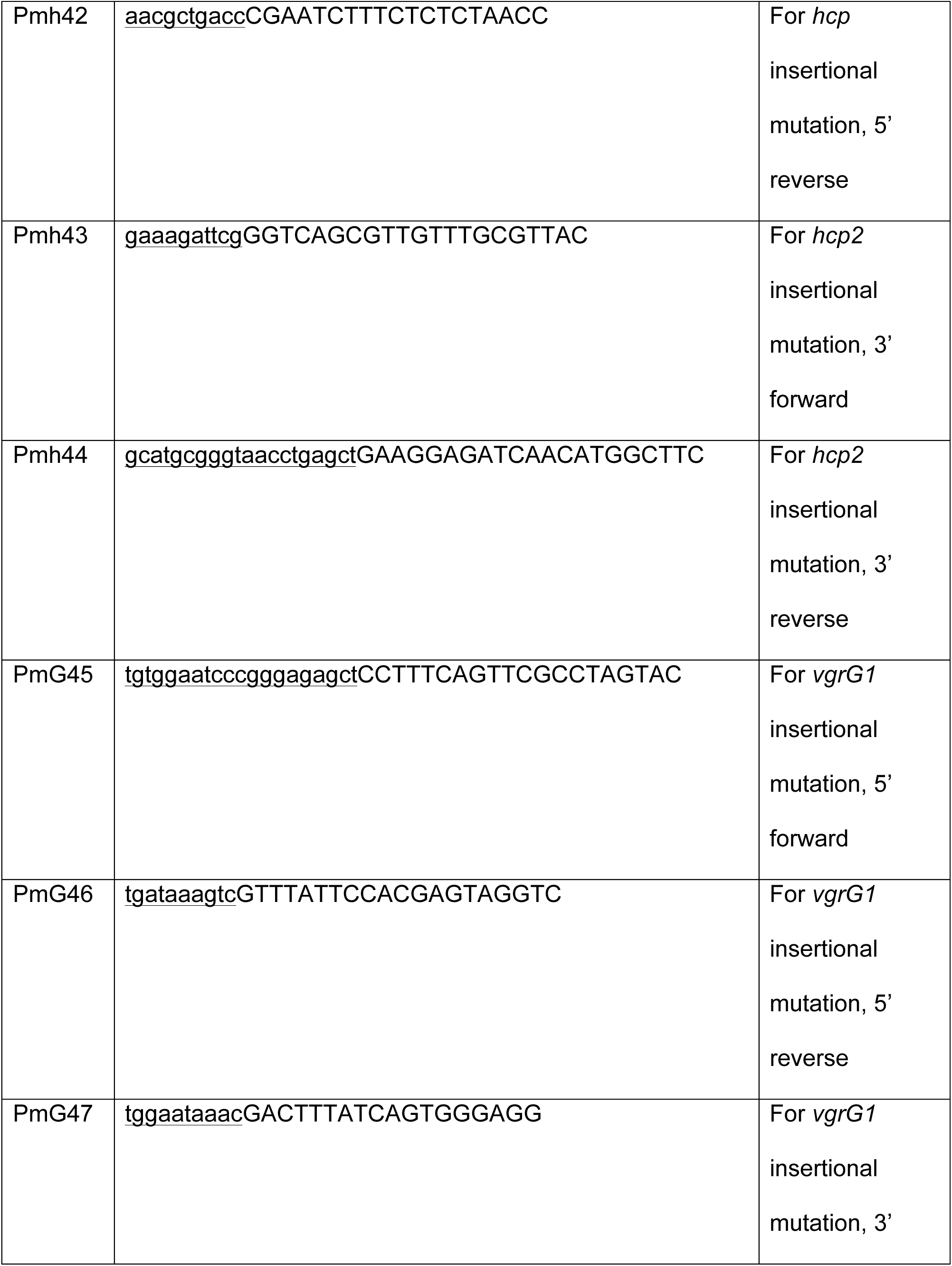
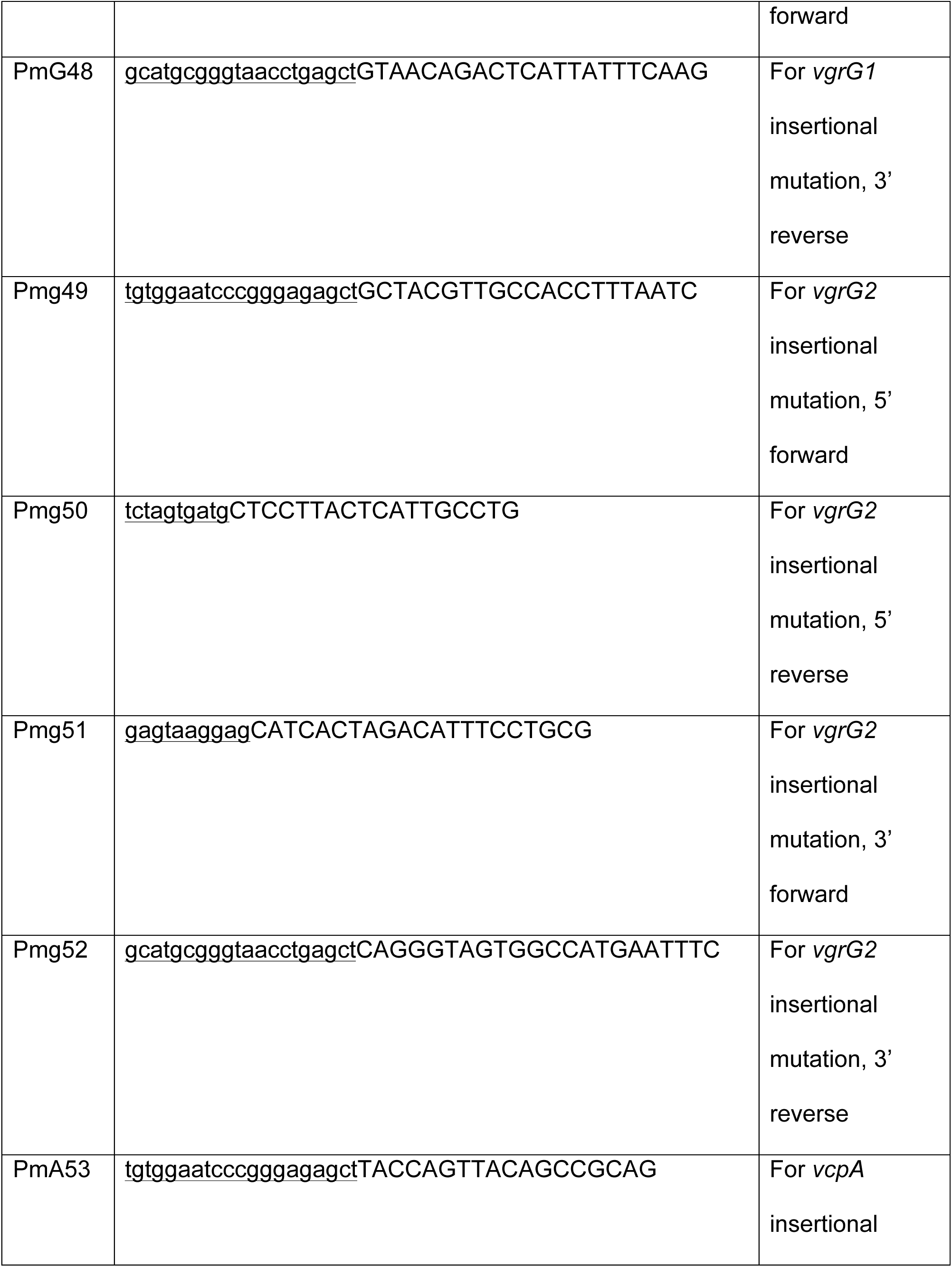
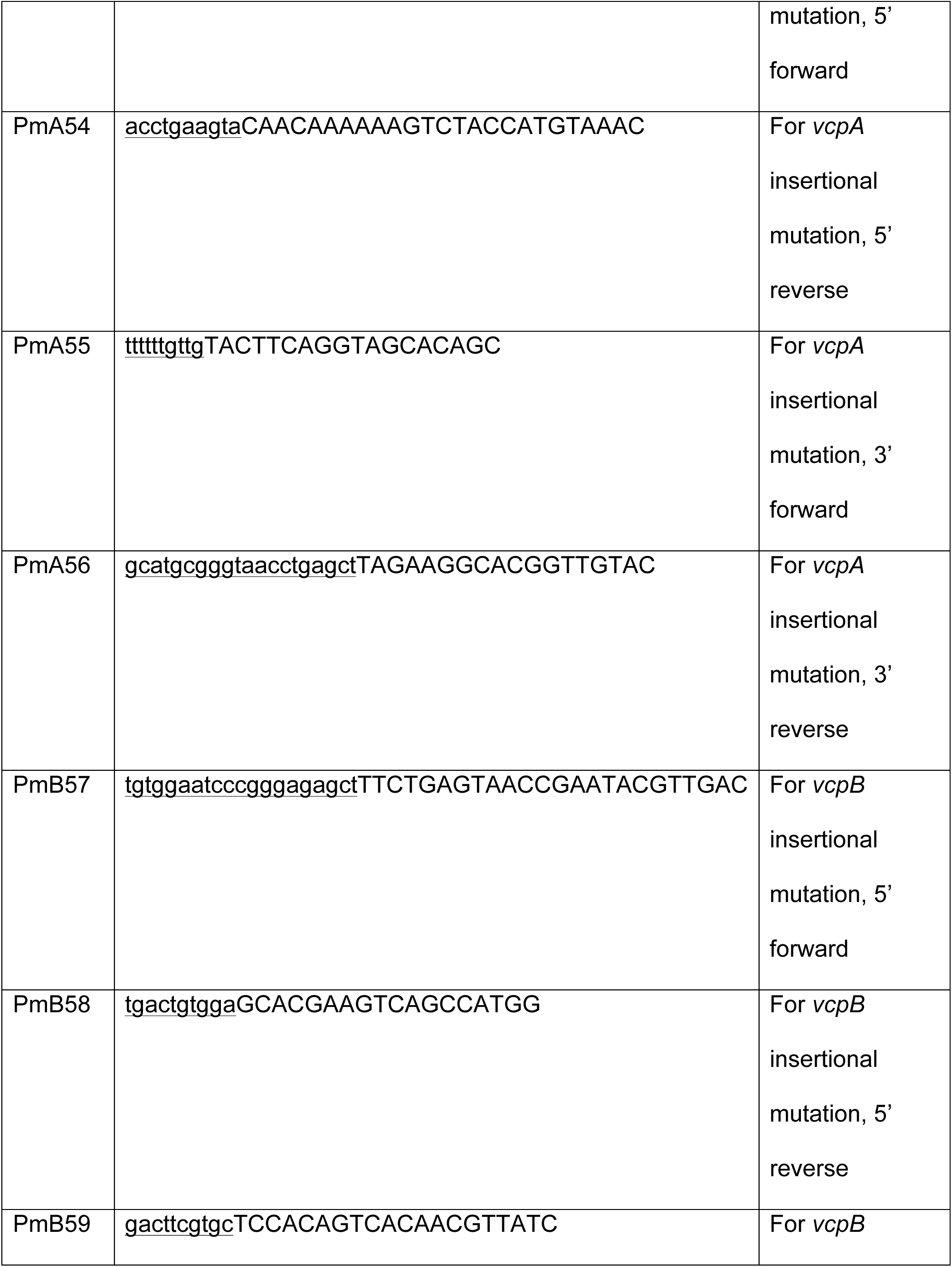
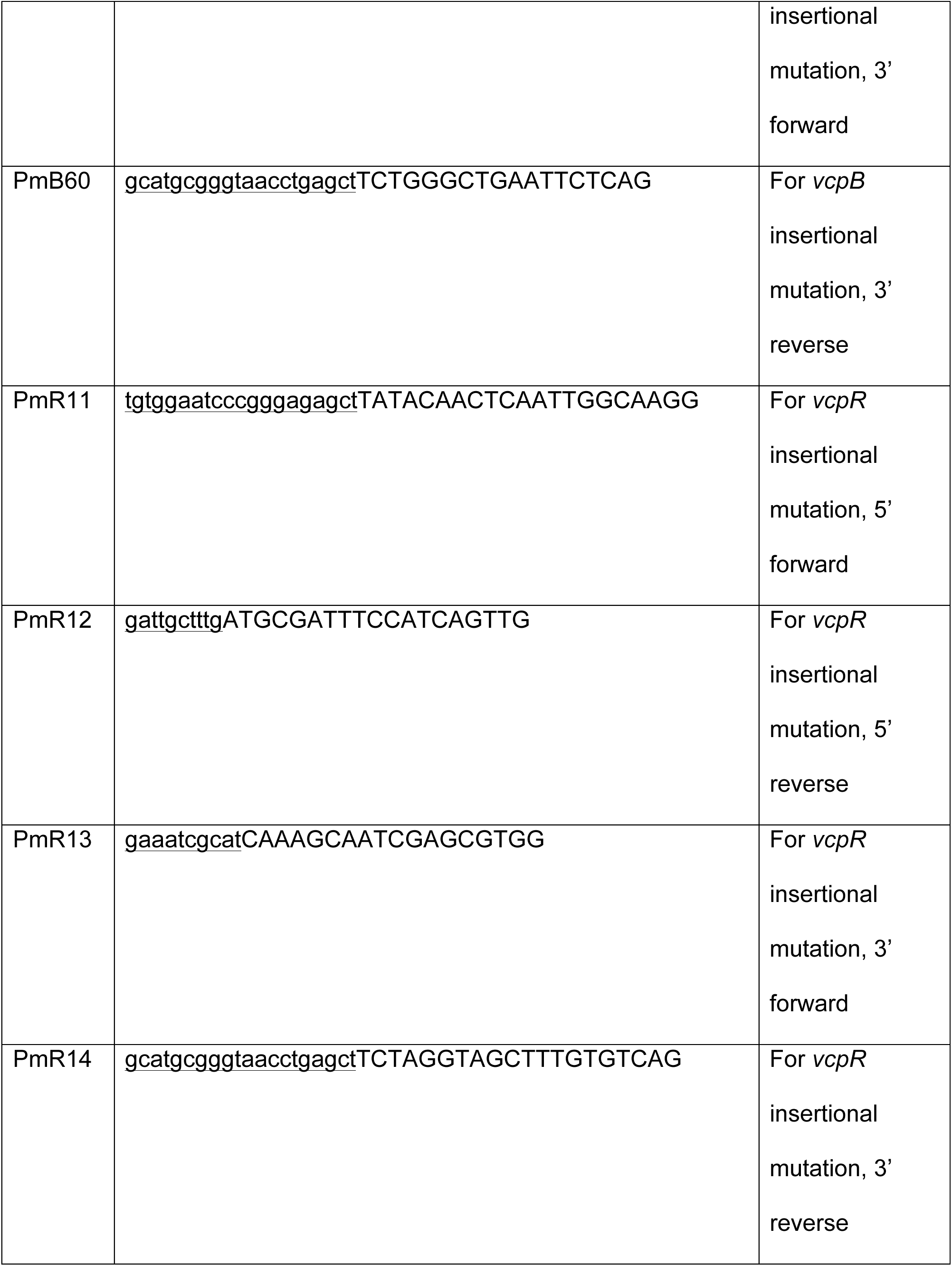
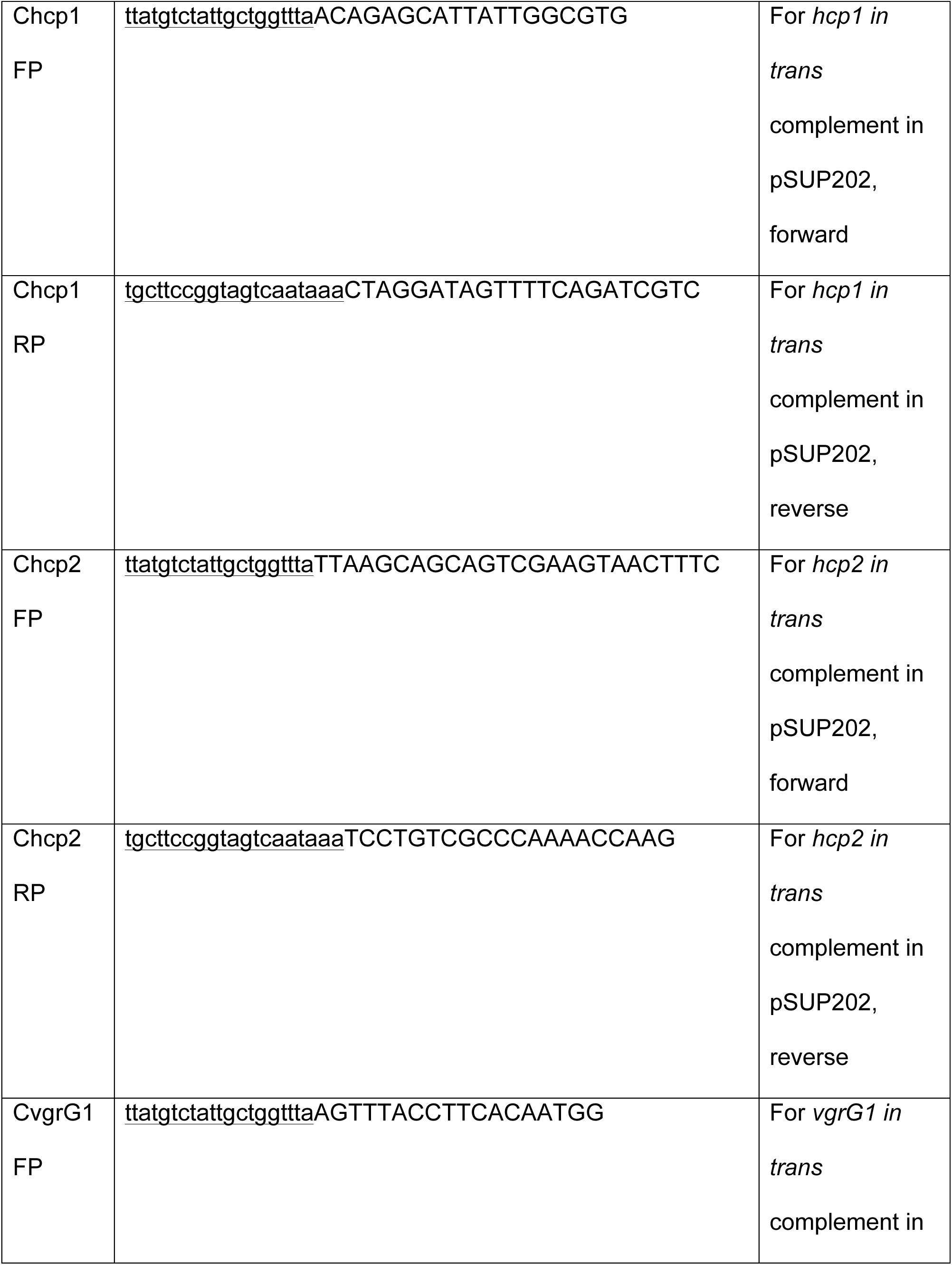
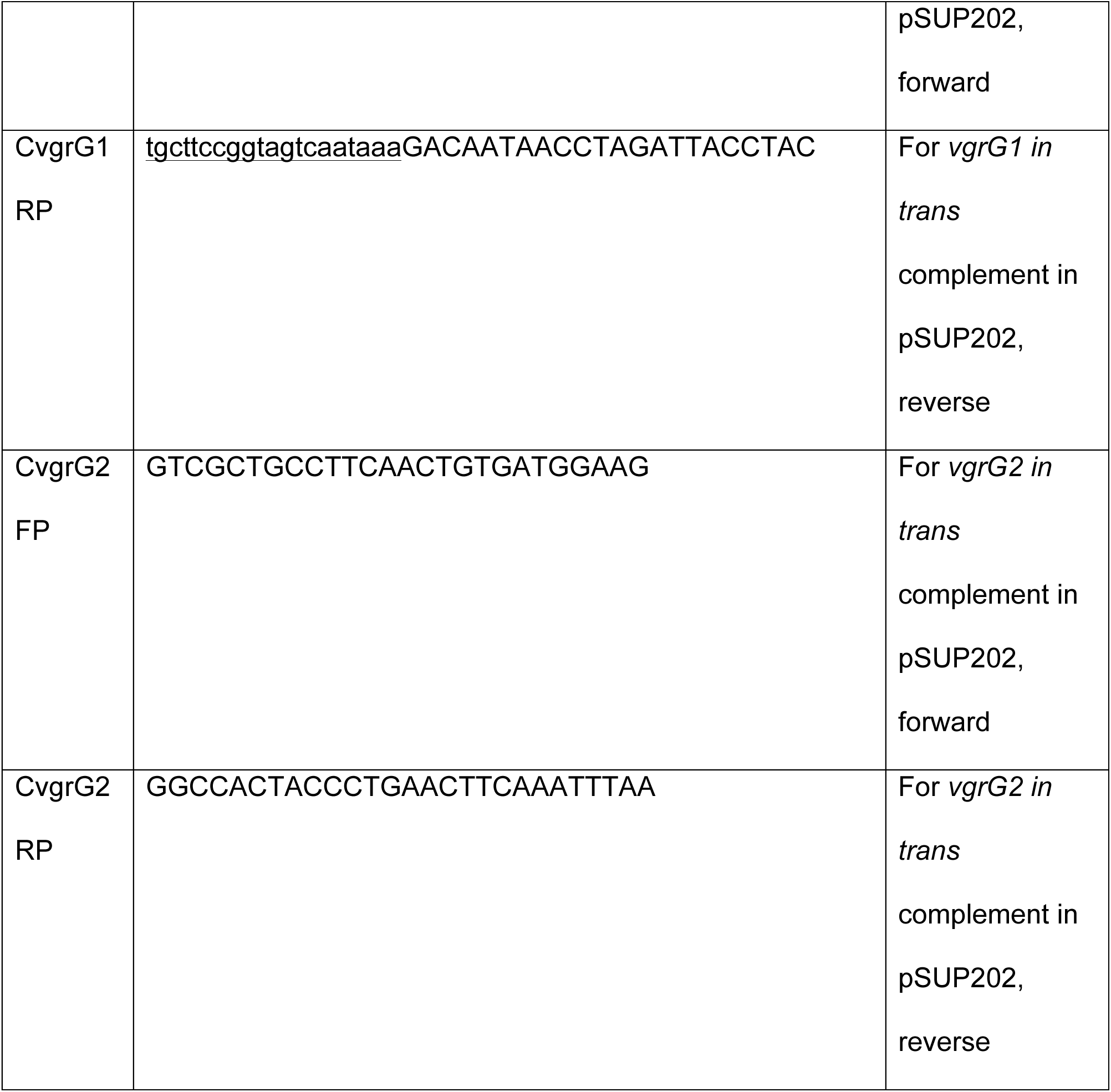
Primers used in this study.

Resolution of the merodiploid state has been previously described by Milton *et al.* (54–56) and subsequently modified. After growth and passage without selection, merodiploid mutants were plated on 3M agar containing 5% sucrose and appropriate antibiotics to select for the double crossover event. The merodiploid mutants were cross-picked onto mYP30Sm^200^ and mYP30Sm^200^Km^80^. Successful growth in the absence of kanamycin indicated a potential allelic exchange and colonies were then screened via PCR for this double crossover event.

### Bacterial killing assays

Assays for determination of T6SS-mediated killing were carried out as described by Salomon *et al.* (17). Briefly, an attacker-to-prey ratio of 4:1 (MOI of 4), based on CFU/ml, was used. A mixture of attacker and prey cells was filtered onto a 0.22 µ filter and placed on appropriate solid growth media for 4 h. The filter was then removed from the agar plate and vortexed for 1 minute in 10 ml NSS, the culture supernatant serially diluted, and plated on appropriate differential media to enumerate the attacker cells and remaining prey cells. TCBS agar was used to select for *Vibrio* spp. and MacConkey agar to select for enteric organisms.

### Larval oyster experimental challenges

Assays for to determine the pathogenicity of *V. coralliilyticus* wild type and mutant strains against eastern oyster larvae were performed as previously described by Zhao *et al.* (2016) with minor modifications. Larval eastern oysters (*Crassostrea virginica*) (6 to 10 days of age, 50 – 150 µm in size) were obtained from the Blount Shellfish Hatchery at Roger William University (Bristol, RI, USA), Virginia Institute of Marine Science (Gloucester Point, VA, USA) or Niantic Bay Shellfish Farm (Niantic, CT, USA), and were allowed to acclimate for 24 h at room temperature with gentle rocking. Next, ∼100 oysters were placed in each well of a 6 well plate containing 5 ml of sterilized filtered artificial seawater at 2.8% salinity. Next, the pathogen, *V. coralliilyticus* RE22Sm (wild type or mutant strain was added to the challenge wells at a final concentration of 10^5^ CFU/ml and incubated for 24 h. Larval oysters were fed with commercial algal paste (20,000 cells/ml; Reed Mariculture Inc., San Jose, CA, USA) in order to promote ingestion of the probiotics. Control wells will include non-treated larvae (with and without pathogen) and larvae incubated with probiotics but not with the pathogen. Each treatment was run in triplicate and each experiment was done at least two times. Larval survival was determined 20-26 h after addition of the pathogen.

The survival rate is calculated using the formula:

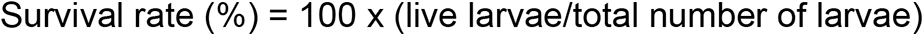

### Statistical analysis

Two-tailed Student’s *t* tests assuming unequal variance were used for all statistical analyses for all detailed experiments. *P* values of < 0.05 were considered to be statistically significant.

## ACKNOWLEDGEMENTS

We thank Ralph Elston for providing *Vibro coralliilyticus* RE22. We also thank Edward Spinard for providing training on multiple techniques used in this study.

This work was supported by a grant from the USDA (2019-67016-29868) to DRN, MGC, and DCR. Funding sources had no role in study design, data collection and interpretation, or the decision to submit the work for publication.

CS & DRN designed the study. CS created the mutant strains used in this study. CS performed all experiments under the supervision of DRN. CS and DRN wrote the manuscript with contributions and edits from DCR and MGC. Manuscript formatting was performed by CS and DRN. All authors read and approved the final version of this manuscript.

## Notes

### Competing Interest Statement

The authors have declared no competing interest.

## References

1. Elston RA, Hasegawa H, Humphrey KI, Polyak IK, Häse CC. 2008. Re-emergence of *Vibrio tubiashii* in bivalve shellfish aquaculture: severity, environmental drivers, geographic extent and management. Dis Aquat Organ 82:119–134.

2. Estes RM, Friedman CS, Elston RA, Herwig RP. 2004. Pathogenicity testing of shellfish hatchery bacterial isolates on Pacific oyster *Crassostrea gigas* larvae. Dis Aquat Organ 58:223–230.

3. Porsby CH, Gram L. 2016. *Phaeobacter inhibens* as biocontrol agent against *Vibrio vulnificus* in oyster models. Food Microbiol 57:63–70.

4. Shruti Chatterjee, Soumya Haldar. 2012. *Vibrio* Related Diseases in Aquaculture and Development of Rapid and Accurate Identification Methods. J Mar Sci Res Dev S1.

5. Spinard E, Kessner L, Gomez-Chiarri M, Rowley DC, Nelson DR. 2015. Draft Genome Sequence of the Marine Pathogen *Vibrio coralliilyticus* RE22. Genome Announc 3:e01432–15.

6. Richards GP, Watson MA, Needleman DS, Church KM, Häse CC. 2015. Mortalities of Eastern and Pacific Oyster Larvae Caused by the Pathogens *Vibrio coralliilyticus* and *Vibrio tubiashii*. Appl Environ Microbiol 81:292–297.

7. Subasinghe R. 2009. Disease control in aquaculture and the responsible use of veterinary drugs and vaccines: the issues, prospects and challenges. Options Méditerranéennes Sér Sémin Méditerranéens 5–11.

8. Culp SJ. 2004. NTP technical report on the toxicity studies of malachite green chloride and leucomalachite green (CAS Nos. 569-64-2 and 129-73-7) administered in feed to F344/N rats and B6C3F1 mice. Toxic Rep Ser 1-F10.

9. K. He X, Wang Z, Nie X, Yang Y, Pan D, Leung AOW, Cheng Z, Yang Y, Li K, Chen K. 2012. Residues of fluoroquinolones in marine aquaculture environment of the Pearl River Delta, South China. Environ Geochem Health 34:323–335.

10. Andersen WC, Turnipseed SB, Roybal JE. 2006. Quantitative and Confirmatory Analyses of Malachite Green and Leucomalachite Green Residues in Fish and Shrimp. J Agric Food Chem 54:4517–4523.

11. Ushijima B, Richards GP, Watson MA, Schubiger CB, Häse CC. 2018. Factors affecting infection of corals and larval oysters by *Vibrio coralliilyticus*. PLOS ONE 13:e0199475.

12. Richards GP, Kingham BF, Shevchenko O, Watson MA, Needleman DS. 2018. Complete Genome Sequence of *Vibrio coralliilyticus* RE22, a Marine Bacterium Pathogenic toward Larval Shellfish. Microbiol Resour Announc 7:e01332–18.

13. Tu KC, Bassler BL. 2007. Multiple small RNAs act additively to integrate sensory information and control quorum sensing in *Vibrio harveyi*. Genes Dev 21:221–233.

14. Guillemette R, Ushijima B, Jalan M, Häse CC, Azam F. 2020. Insight into the resilience and susceptibility of marine bacteria to T6SS attack by *Vibrio cholerae* and *Vibrio coralliilyticus*. PLOS ONE 15:e0227864.

15. Alteri CJ, Mobley HLT. 2016. The Versatile Type VI Secretion System. Microbiol Spectr 4:2.

16. Hachani A, Wood TE, Filloux A. 2016. Type VI secretion and anti-host effectors. Curr Opin Microbiol 29:81–93.

17. Salomon D, Gonzalez H, Updegraff BL, Orth K. 2013. *Vibrio parahaemolyticus* Type VI Secretion System 1 Is Activated in Marine Conditions to Target Bacteria, and Is Differentially Regulated from System 2. PLoS ONE 8:e61086.

18. Ma AT, Mekalanos JJ. 2010. In vivo actin cross-linking induced by *Vibrio cholerae* type VI secretion system is associated with intestinal inflammation. Proc Natl Acad Sci 107:4365–4370.

19. Dubert J, Barja JL, Romalde JL. 2017. New Insights into Pathogenic Vibrios Affecting Bivalves in Hatcheries: Present and Future Prospects. Front Microbiol 8:762.

20. Ma L-S, Hachani A, Lin J-S, Filloux A, Lai E-M. 2014. *Agrobacterium tumefaciens* Deploys a Superfamily of Type VI Secretion DNase Effectors as Weapons for Interbacterial Competition In Planta. Cell Host Microbe 16:94–104.

21. Salmond GPC, Reeves PJ. 1993. Membrance traffic wardens and protein secretion in Gram-negative bacteria. Trends Biochem Sci 18:7–12.

22. Cianfanelli FR, Monlezun L, Coulthurst SJ. 2016. Aim, Load, Fire: The Type VI Secretion System, a Bacterial Nanoweapon. Trends Microbiol 24:51–62.

23. K. Salomon D, Kinch LN, Trudgian DC, Guo X, Klimko JA, Grishin NV, Mirzaei H, Orth K. 2014. Marker for type VI secretion system effectors. Proc Natl Acad Sci 111:9271–9276.

24. Aziz RK, Bartels D, Best AA, DeJongh M, Disz T, Edwards RA, Formsma K, Gerdes S, Glass EM, Kubal M, Meyer F, Olsen GJ, Olson R, Osterman AL, Overbeek RA, McNeil LK, Paarmann D, Paczian T, Parrello B, Pusch GD, Reich C, Stevens R, Vassieva O, Vonstein V, Wilke A, Zagnitko O. 2008. The RAST Server: Rapid Annotations using Subsystems Technology. BMC Genomics 9:75.

25. MacIntyre DL, Miyata ST, Kitaoka M, Pukatzki S. 2010. The *Vibrio cholerae* type VI secretion system displays antimicrobial properties. Proc Natl Acad Sci 107:19520–19524.

26. Shneider MM, Buth SA, Ho BT, Basler M, Mekalanos JJ, Leiman PG. 2013. PAAR- repeat proteins sharpen and diversify the type VI secretion system spike. Nature 500:350–353.

27. Pena RT, Blasco L, Ambroa A, González-Pedrajo B, Fernández-García L, López M, Bleriot I, Bou G, García-Contreras R, Wood TK, Tomás M. 2019. Relationship Between Quorum Sensing and Secretion Systems. Front Microbiol 10:1100.

28. Kimes NE, Grim CJ, Johnson WR, Hasan NA, Tall BD, Kothary MH, Kiss H, Munk AC, Tapia R, Green L, Detter C, Bruce DC, Brettin TS, Colwell RR, Morris PJ. 2012. Temperature regulation of virulence factors in the pathogen *Vibrio coralliilyticus*. ISME J 6:835–846.

29. Karim M, Zhao W, Rowley D, Nelson D, Gomez-Chiarri M. 2013. Probiotic Strains for Shellfish Aquaculture: Protection of Eastern Oyster, *Crassostrea virginica*, Larvae and Juveniles Against Bacterial Challenge. J Shellfish Res 32:401–408.

30. Hasegawa H, Lind EJ, Boin MA, Häse CC. 2008. The Extracellular Metalloprotease of *Vibrio tubiashii* Is a Major Virulence Factor for Pacific Oyster (*Crassostrea gigas*) Larvae. Appl Environ Microbiol 74:4101–4110.

31. Lennings J, West TE, Schwarz S. 2019. The *Burkholderia* Type VI Secretion System 5: Composition, Regulation and Role in Virulence. Front Microbiol 9:3339.

32. Unterweger D, Miyata ST, Bachmann V, Brooks TM, Mullins T, Kostiuk B, Provenzano D, Pukatzki S. 2014. The *Vibrio cholerae* type VI secretion system employs diverse effector modules for intraspecific competition. Nat Commun 5:3549.

33. Ho BT, Dong TG, Mekalanos JJ. 2014. A View to a Kill: The Bacterial Type VI Secretion System. Cell Host Microbe 15:9–21.

34. Kehlet-Delgado H., Häse C.C., Mueller R.S. 2020. Comparative genomic analysis of Vibrios yields insights into genes associated with virulence towards *C. gigas* larvae. BMC Genomics 21:599.

35. Gibbin E, Gavish A, Krueger T, Kramarsky-Winter E, Shapiro O, Guiet R, Jensen L, Vardi A, Meibom A. 2019. *Vibrio coralliilyticus* infection triggers a behavioural response and perturbs nutritional exchange and tissue integrity in a symbiotic coral. ISME J 13:989–1003.

36. Rubio T, Oyanedel D, Labreuche Y, Toulza E, Luo X, Bruto M, Chaparro C, Torres M, de Lorgeril J, Haffner P, Vidal-Dupiol J, Lagorce A, Petton B, Mitta G, Jacq A, Le Roux F, Charrière GM, Destoumieux-Garzón D. 2019. Species-specific mechanisms of cytotoxicity toward immune cells determine the successful outcome of *Vibrio* infections. Proc Natl Acad Sci 116:14238–14247.

37. Grant CE, Bailey TL, Noble WS. 2011. FIMO: scanning for occurrences of a given motif. Bioinformatics 27:1017–1018.

38. Ray A, Schwartz N, de Souza Santos M, Zhang J, Orth K, Salomon D. 2017. Type VI secretion system MIX-effectors carry both antibacterial and anti-eukaryotic activities. EMBO Rep 18:1978–1990.

39. Cianfanelli FR, Diniz JA, Guo M, Cesare VD, Trost M, Coulthurst SJ. 2016. VgrG and PAAR Proteins Define Distinct Versions of a Functional Type VI Secretion System. PLOS Pathog 12:e1005735.

40. Dong TG, Ho BT, Yoder-Himes DR, Mekalanos JJ. 2013. Identification of T6SS- dependent effector and immunity proteins by Tn-seq in *Vibrio cholerae*. Proc Natl Acad Sci 110:2623–2628.

41. Crisan CV, Nichols HL, Wiesenfeld S, Steinbach G, Yunker PJ, Hammer BK. 2021. Glucose confers protection to *Escherichia coli* against contact killing by *Vibrio cholerae*. Sci Rep 11:2935.

42. Tang L, Yue S, Li G-Y, Li J, Wang X-R, Li S-F, Mo Z-L. 2016. Expression, secretion and bactericidal activity of type VI secretion system in *Vibrio anguillarum*. Arch Microbiol 198:751–760.

43. Sana TG, Soscia C, Tonglet CM, Garvis S, Bleves S. 2013. Divergent Control of Two Type VI Secretion Systems by RpoN in Pseudomonas aeruginosa. PLoS ONE 8:e76030.

44. Zhao W, Dao C, Karim M, Gomez-Chiarri M, Rowley D, Nelson DR. 2016. Contributions of tropodithietic acid and biofilm formation to the probiotic activity of *Phaeobacter inhibens*. BMC Microbiol 16:1.

45. Li L, Mou X, Nelson DR. 2013. Characterization of Plp, a phosphatidylcholine-specific phospholipase and hemolysin of *Vibrio anguillarum*. BMC Microbiol 13:271.

46. Coutinho TA, Venter SN. 2009. *Pantoea ananatis*: an unconventional plant pathogen. Mol Plant Pathol 10:325–335.

47. Shyntum DY, Theron J, Venter SN, Moleleki LN, Toth IK, Coutinho TA. 2015. *Pantoea ananatis* Utilizes a Type VI Secretion System for Pathogenesis and Bacterial Competition. Mol Plant Microbe Interact 28:420–431.

48. Zheng J, Leung KY. 2007. Dissection of a type VI secretion system in *Edwardsiella tarda*. Mol Microbiol 66:1192–1206.

49. Pukatzki S, McAuley SB, Miyata ST. 2009. The type VI secretion system: translocation of effectors and effector-domains. Curr Opin Microbiol 12:11–17.

50. Filloux A, Hachani A, Bleves S. 2008. The bacterial type VI secretion machine: yet another player for protein transport across membranes. Microbiology 154:1570–1583.

51. Bingle LE, Bailey CM, Pallen MJ. 2008. Type VI secretion: a beginner’s guide. Curr Opin Microbiol 11:3–8.

52. Gibson DG. 2011. Enzymatic Assembly of Overlapping DNA Fragments. Methods Enzymol 498:349–361.

53. Mou X, Spinard EJ, Driscoll MV, Zhao W, Nelson DR. 2013. H-NS Is a Negative Regulator of the Two Hemolysin/Cytotoxin Gene Clusters in *Vibrio anguillarum*. Infect Immun 81:3566–3576.

54. García-Aljaro C, Melado-Rovira S, Milton DL, Blanch AR. 2012. Quorum-sensing regulates biofilm formation in *Vibrio scophthalmi*. BMC Microbiol 12:287.

55. Milton DL, O’Toole R, Horstedt P, Wolf-Watz H. 1996. Flagellin A is essential for the virulence of *Vibrio anguillarum*. J Bacteriol 178:1310–1319.

56. Newmark KG, O’Reilly EK, Pohlhaus JR, Kreuzer KN. 2005. Genetic analysis of the requirements for SOS induction by nalidixic acid in *Escherichia coli*. Gene 356:69–76.

