## Supplemental Data Tables S1 to S4 for "Two Type VI Secretion Systems in *Vibrio coralliilyticus* RE22Sm exhibit differential target specificity for bacteria prey and oyster larvae"

**Table S1.** T6SS1 genes identified on chromosome 1 (GenBank: CP031472.1) (1)

| Inclusive base pairs | | Strand | Gene | Homologue | Annotation | Putative Function |
|---|---|---|---|---|---|---|
| 675003 | 678314 | - | *tssM* | *vasK, icmF* | IcmF-related protein | Anchoring T6SS to cell wall |
| 678389 | 679591 | - | | *impK, vasF* | Outer membrane protein ImpK/VasF, OmpA/MotB domain | Unknown function |
| 679600 | 680928 | - | *tssK* | *impJ, vasE* | VasE Superfamily | Unknown function |
| 680952 | 681431 | - | *tssJ* | *vasD, lip* | Type VI secretion lipoprotein/VasD | Anchoring T6SS to cell wall |
| 681434 | 682144 | - | *tagH* | *impI* | Uncharacterized protein ImpI/VasC | FHA domain-containing protein, post-translational regulation, putative nuclear signaling domain |
| 682189 | 682338 | - | | | Hypothetical protein | Unknown |
| 682382 | 683101 | - | | | FIG01199604: hypothetical protein | ABC transporter substrate-binding protein [*Vibrio coralliilyticus*]/ SBP bac3 superfamily |
| 683092 | 684090 | - | | | ABC-type uncharacterized transport system, periplasmic component | ABC transporter substrate-binding protein [*Vibrio coralliilyticus*] |
| 684090 | 686621 | - | | *mdtG* | putative inner membrane transport protein | Major Facilitator Superfamily; drug efflux system protein |
| 686659 | 689268 | - | *tssH* | *clpV, vasG* | ClpB protein | ATPase/effector chaperon/recycling TssB/TssC |
| 689346 | 690272 | - | *tssG* | *impH, vasB* | Uncharacterized protein ImpH/VasB | Unknown function/ intracellular trafficking, secretion and vesicular transport |
| 690296 | 692044 | - | *tssF* | *impG, vasA* | Protein ImpG/VasA - phage tail protein needed for Hcp assembly | Unknown function - necessary for Hcp assembly |
| 692040 | 692459 | - | *tssE* | *impF, vasS* | FIG01286925: hypothetical protein | Essential baseplate protein similar to T4 phage gp25 proteins |
| 692462 | 693853 | - | *tssC* | *impC, vipB* | Uncharacterized protein ImpD | Homologous to T4 phage contractile tail sheath proteins |

| | | | | | | |
|---|---|---|---|---|---|---|
| 693918 | 695393 | - | *tssC* | *impC, vipB* | Uncharacterized protein ImpC | Homologous to T4 phage contractile tail sheath proteins |
| 695393 | 695896 | - | *tssB* | *impB, vipA* | Uncharacterized protein ImpB | Homologous to T4 phage contractile tail sheath proteins |
| 695919 | 696437 | - | *tssD* | *hcp1* | Uncharacterized protein ImpD | Effector/Structure: Homologous to T4 phage tube |
| 696475 | 697875 | - | *tssA* | *impA, vasJ* | Uncharacterized protein ImpA | Unknown function - *impA* N terminal domain |
| 698275 | 698421 | - | *paaR* | | PAAR containing protein | Protein with a PAAR motif associated with VgrG piercing structure |
| 698574 | 699140 | - | | | Twin-arginine translocation pathway signal | Hypothetical protein |
| 699148 | 701127 | - | *tssI* | *vgrG1* | VgrG protein | Effector/structure: forms the T6SS piercing structure |
| 701325 | 702128 | - | | | Hypothetical protein | Hypothetical protein |

**Table S2.** T6SS2 genes identified on chromosome 2 (GenBank: CP031473.1) (1)

| Inclusive base pairs | | Strand | Gene | Homologue | Annotation | Putative Function |
|---|---|---|---|---|---|---|
| 87561 | 88349 | - | | *dotU* | Outer membrane protein ImpK/VasF, OmpA/MotB domain | IcmF associated DotU inner membrane anchoring protein |
| 86281 | 87561 | - | *tssK* | *impJ, vasE* | Uncharacterized protein ImpJ/VasE | Unknown function |
| 85778 | 86224 | - | *tssJ* | *vasD, lip* | Type VI secretion lipoprotein/VasD | Anchoring T6SS to cell wall |
| 84349 | 85731 | - | *tagH* | *impI* | Uncharacterized protein ImpI/VasC | FHA domain-containing protein, post-translational regulation |
| 83734 | 84333 | - | | | FIG01199688: hypothetical protein | Outer membrane protein with beta-barrel domain |
| 81901 | 83235 | + | *tssA* | *impA, vasJ* | Uncharacterized protein ImpA | Unknown function - *impA* N terminal domain & DUF1043 superfamily |
| 78503 | 81895 | + | *tssM* | *vasK, icmF* | IcmF-related protein | Anchoring T6SS to cell wall |
| 78013 | 78459 | + | | | Transcriptional regulator, AsnC family | |
| 76943 | 77938 | - | *tssG* | *impH, vasB* | Uncharacterized protein ImpH/VasB | Unknown function |
| 75198 | 76943 | - | *tssF* | *impG, vasA* | Protein ImpG/VasA | Unknown function |
| 74769 | 75176 | - | *tssE* | *impF, vasS* | Uncharacterized protein similar to VCA0109 | Essential baseplate protein similar to T4 phage gp25 proteins |
| 73278 | 74702 | - | *tssC* | *impC, vipB* | Uncharacterized protein ImpC | Homologous to T4 phage contractile tail sheath proteins |
| 72711 | 73217 | - | *tssB* | *impB, vipA* | Uncharacterized protein ImpB | Homologous to T4 phage contractile tail sheath proteins |
| 71118 | 72680 | - | *tssA* | *impA, vasJ* | Uncharacterized protein ImpA | Unknown function |
| 69102 | 71111 | - | *tagE* | *pknA, ppkA* | Serine/threonine protein kinase | Serine/threonine kinase, post-translational regulation |

| | | | | | | |
|---|---|---|---|---|---|---|
| 67955 | 68905 | - | | | FIG01200163: hypothetical protein | No conserved domains |
| 66975 | 67955 | - | | *tagAB* | Pentapeptide repeat family protein | Domain of unknown function |
| 65675 | 66991 | - | | | FIG01200268: hypothetical protein | No conserved domains |
| 65170 | 65652 | - | | | FIG01199591: hypothetical protein | No conserved domains |
| 63092 | 65167 | - | *tssI* | *vgrG2* | VgrG protein | Effector/structure: forms the T6SS piercing structure (potential lysozyme domain) |
| 62502 | 63017 | - | *tssD* | *hcp2* | Hcp protein | Effector/Structure: Homologous to T4 phage tube |
| 59370 | 62039 | + | *tssH* | *clpV* | ClpV protein | MULTISPECIES: ClpV family T6SS ATPase [*Vibrio*] |
| 57746 | 59380 | + | | | Sigma-54 dependent transcriptional regulator | |
| 52921 | 57684 | - | | *tagL* | Hypothetical protein | OmpA family protein [*Vibrio coralliilyticus*] |
| 51663 | 52925 | - | | | Hypothetical protein | No conserved domains |
| 49984 | 51636 | - | | | FIG01201986: hypothetical protein | No conserved domains |
| 48162 | 49121 | + | | *L376_02862* | Cell wall endopeptidase, family M23/M37 | Protein with a peptidase M23 domain, putative endopeptidase effector |

**Table S3. Larval oyster survival after challenge with wild type and mutant strains of *V. coralliilyticus* RE22Sm**

| Treatment[1] | % Survival[2] |
|---|---|
| No Treatment | 91.80% |
| RE22Sm | 48.60% |
| RE22Sm Δ*vcpA* | 70.58% |
| RE22Sm Δ*vcpB* | 72.09% |
| RE22Sm Δ*vcpR* | 80.15% |

[1]Oyster larvae were exposed to RE22Sm wild type and mutant strains ($1\times10^5$ CFU/mL) for 24 h. Oyster larvae treated with artificial seawater served as the negative control.

[2]Larval survival (% ±1 SD) was determined after 24 h challenge. The survival rate is calculated using the formula: Survival rate (%) = 100 x (live larvae/total number of larvae)

**Table S4.** List of core gene and accessory components of the type VI secretion system (T6SS) and putative function derived (2–5)

| Gene | Homologue | Putative Function |
|---|---|---|
| *tssI* | *vgrG* | Effector/structure: forms the T6SS piercing structure |
| *tssD* | *hcp* | Effector/Structure: Homologous to T4 phage tube |
| *tssC* | *impC, vipB* | Homologous to T4 phage contractile tail sheath proteins |
| *tssB* | *impB, vipA* | Homologous to T4 phage contractile tail sheath proteins |
| *tssH* | *clpV, vasG* | ATPase /effector chaperon/recycling TssB/TssC |
| *tssM* | *vasK, icmF* | Anchoring T6SS to cell wall |
| *tssL* | *ompA, dotU* | Anchoring T6SS to cell wall |
| *tssJ* | *vasD, lip* | Anchoring T6SS to cell wall |
| *tssE* | *impF, vasS* | Essential baseplate protein similar toT4 phage gp25 proteins |
| *tssG* | *impH, vasB* | Unknown function |
| *tssF* | *impG, vasA* | Unknown function |
| *tssA* | *impA, vasJ* | Unknown function |
| *tssK* | *impJ, vasE* | Unknown function |
| *tagB* | *BB0796* | Protein with a pentapeptide_4 domain, unknown function |
| *tagAB* | *BB0795* | Protein with a pentapeptide_4 domain, unknown function |
| *tagE* | *pknA/ppkA* | Serine/threonine kinase, post-translational regulation |
| *tagF* | *impM, sciT* | Unknown function |
| *tagG* | *pppA* | Serine/threonine phosphatase, post-translational regulation |
| *tagH* | *impI* | FHA domain-containing protein, post-translational regulation |
| *tagJ* | *impE* | Unknown function |
| *tagL* | *c3389* | Protein with an OmpA C-like domain, unknown function |
| | *VCA0105* | Protein with a PAAR-motif associated with VgrG piercing structure |
| | *L376_02862* | Protein with a peptidase M_23 domain, putative endopeptidase effector |
| | *Ebc_4130* | Protein with an esterase-lipase domain, unknown function |

References:


1. Richards GP, Kingham BF, Shevchenko O, Watson MA, Needleman DS. 2018. Complete Genome Sequence of *Vibrio coralliilyticus* RE22, a Marine Bacterium Pathogenic toward Larval Shellfish. Microbiol Resour Announc 7:e01332-18. doi:10.1128/MRA.01332-18

2. Pukatzki S, Ma AT, Sturtevant D, Krastins B, Sarracino D, Nelson WC, Heidelberg JF, Mekalanos JJ. 2006. Identification of a conserved bacterial protein secretion system in *Vibrio cholerae* using the *Dictyostelium* host model system. Proc Natl Acad Sci 103:1528–1533.

3. Zheng J, Leung KY. 2007. Dissection of a type VI secretion system in *Edwardsiella tarda*. Mol Microbiol 66:1192–1206.

4. Filloux A, Hachani A, Bleves S. 2008. The bacterial type VI secretion machine: yet another player for protein transport across membranes. Microbiology 154:1570–1583.

5. Bingle LE, Bailey CM, Pallen MJ. 2008. Type VI secretion: a beginner’s guide. Curr Opin Microbiol 11:3–8.
